# SpatialDEG: Identification of differentially expressed genes by leveraging spatial information in spatially resolved transcriptomic studies

**DOI:** 10.1101/2022.05.10.491324

**Authors:** Yi Yang, Jeffrey ChunTatt Lim, Cedric Chuan Young Ng, Jing Yi Lee, Joe Yeong, Lei Sun, Jin Liu

## Abstract

**Motivation:** Spatially resolved transcriptomics (SRT) technologies have been developed to simultaneously profile gene expression while retaining physical information. To explore differentially expressed genes using SRT in the context of various conditions, statistical methods are needed to perform spatial differential expression analysis.

**Results:** We propose that a new probabilistic framework, spatialDEG, can perform differential expression analysis by leveraging spatial information on gene expression with spatial information. SpatialDEG utilizes the average information algorithm and can be scalable to tens of thousands of genes. Comprehensive simulations demonstrated that spatialDEG can identify genes differentially expressed in tissues across different conditions with a controlled type-I error rate. We further applied spatialDEG to analyze datasets for human dorsolateral prefrontal cortex and mouse whole liver.

**Availability:** The R package spatialDEG can be downloaded from https://github.com/Shufeyangyi2015310117/spatialDEG.

## 1 Introduction

Spatially resolved transcriptomics (SRT) technologies, which encompass a set of newly developed techniques to simultaneously characterize gene expression profiles at multiple tissue locations while retaining information on the physical locations, have provided unprecedented opportunities to study the spatial distribution of cell types within tissues [13, 20, 12]. When SRT datasets are collected from samples under different conditions, a natural question arises as how to identify the genes differentially expressed under the different conditions.

Many statistical methods have been developed to perform differential expression analysis of single-cell (sc)RNA-seq datasets from different conditions when no spatial information is available. These methods can be broadly categorized into two groups. The first group involves methods developed for bulk RNA-seq data that can be also applied to scRNA-seq data, e.g., DESeq [1], edgeR [18], EBSeq [11], baySeq [5], NBPSeq [3], Limma [17], etc. The second group of methods, e.g., MAST [4], scDD [10], EMDomics [15], D3E [2], SAMstrt [8], DEsingle [14], etc., were developed to account for the unique features in scRNA-seq data. However, all these methods were developed to perform differential expression analysis of either bulk RNA-seq or scRNA-seq data across different conditions while ignoring spatial variation in the tissues.

In this note, we introduce the tool spatialDEG, which is able to perform spatial differential expression analysis. SpatialDEG partitions expression variations into spatial and non-spatial components and identifies the expression variation associated with each spatial location. Using a likelihood ratio test, spatialDEG identifies genes differentially expressed under different conditions. The computational cost of spatialDEG is O(*n*^2^), and spatialDEG is scalable to large sample sizes. We first applied spatialDEG to analyze 10x Genomics Visium datasets from human dorsolateral prefrontal cortex samples as negative controls and then applied spatialDEG to analyze 10x Genomics Visium datasets from mouse whole liver. The results showed that SpatialDEG effectively controlled type-I error rate while identifying differentially expressed genes and accounting for spatial variation.

## 2 Results

### 2.1 Simulation results

In the simulation studies, we focused on two scenarios: (a) an assessment of the type-I error rate under the null hypothesis, and (b) an assessment of the power under the alternative hypothesis.

In Scenario (a), we discovered that the type-I error rate was well controlled under different numbers of spots, heritability, and ground truth kernels (Supplementary Figure 1-24). In Scenario (b), we found there was an increase in power as the difference in the effects under the two conditions increased.

### 2.2 Real data analysis

In the real data analysis, we analyzed two datasets, one from human dorsolateral prefrontal cortex (DLPFC) and one from mouse whole liver tissue, both of which were sequenced on a 10x Genomics Visium platform. To gather the human DLPFC data, a total of 12 samples from three individuals (four samples from each individual) were studied. In our analysis, we evaluated six pairs of samples from the same individual, and there were no differences in the gene expression patterns. The qq-plots, which are shown in Supplementary Figure 33, demonstrate that the type-I error rate was well controlled for all six pairs.

In the datasets for mouse whole liver, four samples were collected from four mice, with two mice kept under one of two conditions, i.e., wild type (WT) or high-fat diet (HFD). We applied spatialDEG to identify the differentially expressed genes between the WT and HFD pairs. The qq-plots for these four pairs are provided in Fig 1. Note that *p*-values less than 1e-10 were truncated to 1e-10 in the qq-plots to save space. Using spatialDEG, we identified 33, 66, 40, and 86 genes that were spatially differentially expressed for the pairs HFD1 vs WT1, HFD1 vs WT2, HFD3 vs WT1, and HFD3 vs WT2, respectively, at an FDR of 5%. Details of these genes are provided in Supplementary Table 4 and the supplementary Excel file.

**Figure 1:**
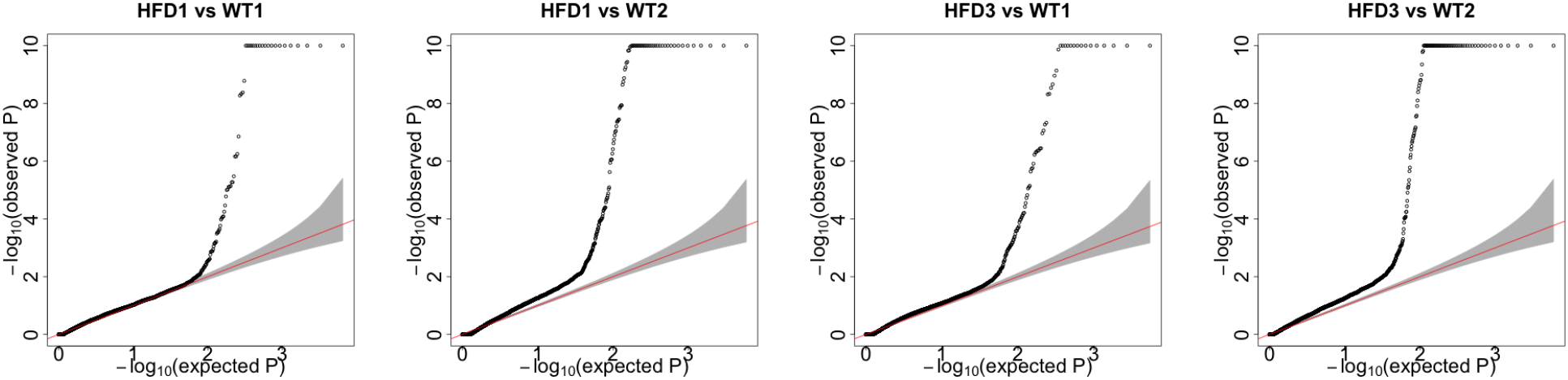
qq-plots of the p-values generated by spatialDEG from the real dataset for mouse whole liver for the pairs HFD1 vs WT1, HFD1 vs WT2, HFD3 vs WT1, and HFD3 vs WT2

We carefully examined the genes that are spatially differentially expressed. As shown in Supplementary Table 4, the pairs HFD1 vs WT1, HFD1 vs WT2, HFD3 vs WT1, and HFD3 vs WT2, were reported to have 10, 17, 16, and 27 differentially expressed genes, respectively, in a previous publication [16]; similarly, the pairs HFD1 vs WT1, HFD1 vs WT2, HFD3 vs WT1, and HFD3 vs WT2 were previously reported to have 8, 28, 15, and 34 differentially expressed genes, respectively, in the published literature [9]. Among the genes that were spatially differentially expressed, *Mup1*, which is a regulator of glucose and lipid metabolism in the mouse [21], was identified in both pairs HFD1 vs WT1 and HFD3 vs WT1. Similarly, *CYP2E1*, identified in both HFD1 vs WT1 and HFD3 vs WT1, belongs to the cytochrome P450 family, is involved in xenobiotic metabolism, and plays an important role during catabolic processes [7, 6]. Additionally, the higher expression levels of *CYP2A5* in HFD-fed mice are in agreement with the pattern described in a previous report [19].

## 3 Discussion

We developed a tool, spatialDEG, to perform differential expression analysis in the spatial context of tissues. After obtaining low-dimensional embeddings for spots (using either principal component analysis or DR-SC [12]), spatialDEG was used to identify genes differentially expressed in tissues across different conditions while accounting for biological differences among cell types. We applied spatialDEG to two 10x Genomics Visium datasets and showed that spatialDEG not only controlled type-I error rates but also identified spatial genes differentially expressed across different conditions.

