## supplement for "SpatialDEG: Identification of differentially expressed genes by leveraging spatial information in spatially resolved transcriptomic studies"

### 1 MATERIALS AND METHODS

#### 1.1 Model setting

Suppose we have two gene expression profiles, $\mathbf{y}_1 = [y_{11}, \cdots, y_{1n_1}]^\top$ and $\mathbf{y}_2 = [y_{21}, \cdots, y_{2n_2}]^\top$, measured at spots $n_1$ and $n_2$, respectively, generated in different experimental conditions. For the simplicity of representation, let $\mathbf{y} = [\mathbf{y}_1^\top, \mathbf{y}_2^\top]^\top$ denote the concatenated gene expression profiles of $\mathbf{y}_1$ and $\mathbf{y}_2$. We also have an $n$-by-$q$ matrix of covariates $\mathbf{W} \in \mathbb{R}^{n\times q}$ ($n = n_1 + n_2$), whose first column is a vector of ones corresponding to the intercept. To detect the difference between the two gene expressions $\mathbf{y}_1$ and $\mathbf{y}_2$, we construct a vector of indicators $\mathbf{z} = [z_1, \cdots, z_n]^\top$ whose entries take a value of 0 in one experimental condition or 1 in another experimental condition. In addition, we are able to obtain the spatial coordinates of all spots in these two gene expressions. Let $\mathbf{S}_1 = [\mathbf{s}_1, \cdots, \mathbf{s}_{n_1}] \in \mathbb{R}^{p\times n_1}$ and $\mathbf{S}_2 = [\mathbf{s}_1, \cdots, \mathbf{s}_{n_2}] \in \mathbb{R}^{p\times n_2}$ denote the matrix of spatial coordinates for the two gene expressions $\mathbf{y}_1$ and $\mathbf{y}_2$, respectively. Note that the dimension of coordinates often depends on a specific spatial transcriptomics technique. For example, $p$ is equal to 2 in 10x Genomics Visium data. Based on these spatial coordinates, we can construct two types of kernels, a Gaussian kernel and a Cosine kernel, which are often adopted to characterize spatial variation in spatially resolved transcriptomic studies. Then we model the concatenated gene expression $\mathbf{y}$ with the following linear model:

$$\begin{aligned} &\mathbf{y} = \mathbf{W}\boldsymbol{\beta} + \mathbf{z}\mu + \mathbf{e} \\ &\mathbf{e} \sim \mathcal{N}\left(\mathbf{0}, \begin{bmatrix} \sigma_1^2\mathbf{K}_1 & \mathbf{0} \\ \mathbf{0} & \sigma_2^2\mathbf{K}_2 \end{bmatrix} + \sigma_e^2\mathbf{I}\right) \end{aligned} \tag{1}$$

where $\mathcal{N}(\mathbf{r}|\mathbf{m}, \boldsymbol{\Sigma})$ denotes the Gaussian distribution with mean $\mathbf{m}$ and covariance matrix $\boldsymbol{\Sigma}$. $\boldsymbol{\beta} = [\beta_1, \cdots, \beta_q]^\top$ is a vector of effect size corresponding to the covariates. $\mu$ is the effect size of the indicators and can reflect the difference between the two gene expressions $\mathbf{y}_1$ and $\mathbf{y}_2$; therefore, we will test it later. To account for both spatial and non-spatial variation in the data, we consider a more accurate random error term, $\mathbf{e}$, as showed in equation (1). In general, $\mathbf{e}$ consists of two components, a spatial component to accommodate spatial variation in gene expression and a non-spatial component to accommodate the usual non-spatial variation in gene expression. In the spatial component, we employ a block diagonal matrix with two kernel functions, $\mathbf{K}_1$ and $\mathbf{K}_2$, to characterize the spatial variation in gene expression $\mathbf{y}_1$ and $\mathbf{y}_2$, respectively; while, in the non-spatial component, we employ the usual $n$-by-$n$ identity matrix $\mathbf{I}$. Obviously, $\sigma_1^2$ and $\sigma_2^2$ are the magnitudes of the expression variances associated with kernels $\mathbf{K}_1$ and $\mathbf{K}_2$, respectively, and $\sigma_e^2$ is the magnitude of the variance associated with $\mathbf{I}$. They depict the proportions of both the spatial and the non-spatial variation. In this part, we let $\boldsymbol{\theta} = \{\sigma_1^2, \sigma_2^2, \sigma_e^2, \boldsymbol{\beta}, \mu\}$ denote the collection of parameters in the model.

##### 1.2 Covariance Kernel Functions

Similar to SPARK [10], we consider two common types of kernel functions, Gaussian and Cosine, to model the gene expression variation associated with spatial location. The Gaussian kernel function is defined as

$$\mathbf{K}_g(\mathbf{s}_i, \mathbf{s}_j) = \exp\left(-\frac{\|\mathbf{s}_i - \mathbf{s}_j\|^2}{2l^2}\right) \tag{2}$$

where $\|\mathbf{s}_i - \mathbf{s}_j\| = \sqrt{\sum_{k=1}^{p}(\mathbf{s}_{ik} - \mathbf{s}_{jk})^2}$ is the Euclidean distance, and $l$ is the length scale parameters that are used to characterize the size of the focal expression patterns. The Cosine kernel function is defined as

$$\mathbf{K}_c(\mathbf{s}_i, \mathbf{s}_j) = \cos\left(2\pi\frac{\|\mathbf{s}_i - \mathbf{s}_j\|}{\phi}\right) \tag{3}$$

where $\phi$ is the periodicity parameters that are used to characterize the frequency of the periodic expression pattern. In this paper, we called both the length scale parameter $l$ and the periodicity parameter $\phi$ the kernel-related parameters.

##### 1.3 Average Information algorithm

The log-likelihood for the model (1) can be written as

$$\mathcal{L}(\boldsymbol{\theta}) = -\frac{n}{2}\log(2\pi) - \frac{1}{2}\log|\mathbf{P}^{-1}| - \frac{1}{2}(\mathbf{y} - \mathbf{W}\boldsymbol{\beta} - \mathbf{z}\mu)^\top\mathbf{P}(\mathbf{y} - \mathbf{W}\boldsymbol{\beta} - \mathbf{z}\mu) \tag{4}$$

where the precision matrix is $\mathbf{P} = \left[\begin{bmatrix} \sigma_1^2\mathbf{K}_1 & \mathbf{0} \\ \mathbf{0} & \sigma_2^2\mathbf{K}_2 \end{bmatrix} + \sigma_e^2\mathbf{I}\right]^{-1}$ and $|\mathbf{P}|$ denotes the determinant of matrix $\mathbf{P}$. To efficiently estimate the parameters $\boldsymbol{\theta}$, we borrow the idea of [4] to factor equation (4) with the Eigen decomposition of kernels $\mathbf{K}_i = \mathbf{U}_i\mathbf{D}_i\mathbf{U}_i^\top (i = 1, 2)$. Then equation (4) can be factored into the following form:

$$\mathcal{L}(\boldsymbol{\theta}) = -\frac{n}{2}\log(2\pi) - \frac{1}{2}\log|\widetilde{\mathbf{P}}^{-1}| - \frac{1}{2}(\mathbf{U}^\top\mathbf{y} - \mathbf{U}^\top\mathbf{W}\boldsymbol{\beta} - \mathbf{U}^\top\mathbf{z}\mu)^\top\widetilde{\mathbf{P}}(\mathbf{U}^\top\mathbf{y} - \mathbf{U}^\top\mathbf{W}\boldsymbol{\beta} - \mathbf{U}^\top\mathbf{z}\mu) \tag{5}$$

where the new notations are $\widetilde{\mathbf{P}} = \mathbf{U}^\top \mathbf{P}\mathbf{U}$, $\mathbf{U} = \begin{bmatrix} \mathbf{U}_1 & \mathbf{0} \\ \mathbf{0} & \mathbf{U}_2 \end{bmatrix}$. Thus, we are capable of precalculating the Eigen decomposition of matrix $\mathbf{K}_i$ at the beginning of the algorithm and avoid the matrix-matrix multiplication in each iteration, which will hugely improve the efficiency of the algorithm.

Next, we infer the model parameters $\boldsymbol{\theta}$ and kernel-related parameters $l$ or $\phi$ based on the log-likelihood (5). First, all the parameters can be classified into three classes based on the method of estimation. The first class comprises the parameters for fixed effects $\boldsymbol{\beta}$ and $\mu$, and we can derive the corresponding closed form easily as follows:

$$\begin{aligned} \boldsymbol{\beta} &= (\mathbf{U}\mathbf{W}^\top \widetilde{\mathbf{P}} \mathbf{U}^\top \mathbf{W})^{-1} \mathbf{U}\mathbf{W}^\top \widetilde{\mathbf{P}} (\mathbf{U}^\top \mathbf{y} - \mathbf{U}^\top \mathbf{z}\mu) \\ \mu &= (\mathbf{U}\mathbf{z}^\top \widetilde{\mathbf{P}} \mathbf{U}^\top \mathbf{z})^{-1} \mathbf{U}\mathbf{z}^\top \widetilde{\mathbf{P}} (\mathbf{U}^\top \mathbf{y} - \mathbf{U}^\top \mathbf{W}\boldsymbol{\beta}) \end{aligned} \tag{6}$$

It is apparent that the estimation of these parameters is very easy and efficient. The second class comprises the error-related parameters $\boldsymbol{\theta}_1 = \{\sigma_1^2, \sigma_2^2, \sigma_e^2\}$. Theoretically, there are no closed forms of solutions for them. Therefore, we have to adopt the Average Information (AI) algorithm [1]. In general, the main idea of the AI algorithm is to first calculate the observed information of the log-likelihood (5) with respect to $\boldsymbol{\theta}_1$ and its expectation; then take the average of the above observed and expected information, and finally, update the error-related parameters with the following Newton-Raphson formula:

$$\boldsymbol{\theta}_1^{t+1} = \boldsymbol{\theta}_1^t - \mathrm{AI}^{-1} \frac{\partial \mathcal{L}(\boldsymbol{\theta}_1)}{\partial \boldsymbol{\theta}_1} \tag{7}$$

where the notation AI denotes the AI matrix and $\frac{\partial \mathcal{L}(\boldsymbol{\theta}_1)}{\boldsymbol{\theta}_1}$ is the first order derivative of the log-likelihood (5) with respect to $\boldsymbol{\theta}_1$. The details of the derivation of the AI matrix are in the section 1.5 of the supplementary materials. The third class of parameters comparises the length scale or the periodicity parameters, and we can search for the optimal value by maximizing the log-likelihood (5) on a grid of predefined length scale $l_1, \cdots, l_{k_1}$and the periodicity parameter $\phi_1, \cdots, \phi_{k_2}$to select the best optimal kernel functions:

$$\widehat{\phi} = \underset{\phi \in \{l_1, \cdots, l_{k_1}, \phi_1, \cdots, \phi_{k_2}\}}{\arg\max} \mathcal{L}(\phi) \tag{8}$$

Where $k_1$ and $k_2$ denote the number of the length scales and periodicity parameters, respectively. Note that $k_1$ and $k_2$ can be specified by users in the software. Lastly, we summarize the whole algorithm as Algorithm 1 in the section 1.6 of the supplementary materials.

#### 1.4 Hypothesis testing

We then aim to identify the differentially expressed genes in the spatially resolved transcriptome studies. In our model, it is equivalent to testing

$$\mathcal{H}_0 : \quad \mu = 0 \qquad \text{vs} \qquad \mathcal{H}_1 : \quad \mu \neq 0 \tag{9}$$

Here, we perform the log-likelihood ratio test. The test statistics can be expressed as

$$\lambda = 2\left(\mathcal{L}(\widehat{\boldsymbol{\theta}}_{\mathcal{H}_1}) - \mathcal{L}(\widehat{\boldsymbol{\theta}}_{\mathcal{H}_0})\right) \tag{10}$$

where $\mathcal{L}(\widehat{\boldsymbol{\theta}}_{\mathcal{H}_1})$ and $\mathcal{L}(\widehat{\boldsymbol{\theta}}_{\mathcal{H}_0})$ denote the log-likelihood with the estimated parameters in the alternative and null hypotheses, respectively. Under the null hypothesis, the statistics follows a chi-square distribution with one degree of freedom [11].

#### 1.5 Details of derivation in the average information algorithm

In this section, we describe how to derive the AI algorithm in detail. First, for ease of notation, we let

$$\widetilde{\mathbf{W}} = \mathbf{U}^\top \mathbf{W} \quad \widetilde{\mathbf{y}} = \mathbf{U}^\top \mathbf{y} \quad \widetilde{\mathbf{z}} = \mathbf{U}^\top \mathbf{z} \tag{11}$$

Thus, the log-likelihood for model (1) in the section 1.1 can be written as

$$\mathcal{L}(\boldsymbol{\theta}) = -\frac{n}{2}\log(2\pi) - \frac{1}{2}\log|\widetilde{\mathbf{P}}^{-1}| - \frac{1}{2}(\widetilde{\mathbf{y}} - \widetilde{\mathbf{W}}\boldsymbol{\beta} - \widetilde{\mathbf{z}}\mu)^\top \widetilde{\mathbf{P}}(\widetilde{\mathbf{y}} - \widetilde{\mathbf{W}}\boldsymbol{\beta} - \widetilde{\mathbf{z}}\mu) \tag{12}$$

where

$$\widetilde{\mathbf{P}} = \left[\begin{bmatrix} \sigma_1^2\mathbf{D}_1 & \mathbf{0} \\ \mathbf{0} & \sigma_2^2\mathbf{D}_2 \end{bmatrix} + \sigma_e^2\mathbf{I}\right]^{-1} \tag{13}$$

Note that $\widetilde{\mathbf{P}}$ becomes a diagonal matrix, which facilitates the calculation of the AI matrix. Prior to the calculation of the first derivative, we require the following identities.

Assuming $\mathbf{U} = \mathbf{U}(x)$, then the following identities hold:

$$\frac{\partial \ln|\mathbf{U}|}{\partial x} = \mathrm{Tr}\left(\mathbf{U}^{-1}\frac{\partial \mathbf{U}}{\partial x}\right) \tag{14}$$

$$\frac{\partial \mathbf{U}^{-1}}{\partial x} = -\mathbf{U}^{-1}\frac{\partial \mathbf{U}}{\partial x}\mathbf{U}^{-1} \tag{15}$$

We can then obtain the first order derivatives of the log-likelihood with respect to the parameters $\{\sigma_1^2, \sigma_2^2, \sigma_e^2\}$

$$\begin{aligned}
\frac{\partial \mathcal{L}}{\partial \sigma_e^2} &= \frac{1}{2}(\widetilde{\mathbf{y}} - \widetilde{\mathbf{W}}\boldsymbol{\beta} - \widetilde{\mathbf{z}}\mu)^\top \widetilde{\mathbf{P}}\widetilde{\mathbf{P}}(\widetilde{\mathbf{y}} - \widetilde{\mathbf{W}}\boldsymbol{\beta} - \widetilde{\mathbf{z}}\mu) - \frac{1}{2}\mathrm{Tr}(\widetilde{\mathbf{P}}) \\
\frac{\partial \mathcal{L}}{\partial \sigma_1^2} &= \frac{1}{2}(\widetilde{\mathbf{y}} - \widetilde{\mathbf{W}}\boldsymbol{\beta} - \widetilde{\mathbf{z}}\mu)^\top \widetilde{\mathbf{P}}\mathbf{S}_1\widetilde{\mathbf{P}}(\widetilde{\mathbf{y}} - \widetilde{\mathbf{W}}\boldsymbol{\beta} - \widetilde{\mathbf{z}}\mu) - \frac{1}{2}\mathrm{Tr}(\widetilde{\mathbf{P}}\mathbf{S}_1) \\
\frac{\partial \mathcal{L}}{\partial \sigma_2^2} &= \frac{1}{2}(\widetilde{\mathbf{y}} - \widetilde{\mathbf{W}}\boldsymbol{\beta} - \widetilde{\mathbf{z}}\mu)^\top \widetilde{\mathbf{P}}\mathbf{S}_2\widetilde{\mathbf{P}}(\widetilde{\mathbf{y}} - \widetilde{\mathbf{W}}\boldsymbol{\beta} - \widetilde{\mathbf{z}}\mu) - \frac{1}{2}\mathrm{Tr}(\widetilde{\mathbf{P}}\mathbf{S}_2)
\end{aligned} \tag{16}$$

where $\mathbf{S}_1$ and $\mathbf{S}_2$ are defined as follows:

$$\mathbf{S}_1 = \begin{bmatrix} \mathbf{D}_1 & \mathbf{0} \\ \mathbf{0} & \mathbf{0} \end{bmatrix} \quad \mathbf{S}_2 = \begin{bmatrix} \mathbf{0} & \mathbf{0} \\ \mathbf{0} & \mathbf{D}_2 \end{bmatrix} \tag{17}$$

Similarly, referring to the matrix calculus in Wikipedia, the following identities hold:

$$d(\mathrm{Tr}(\mathbf{X})) = \mathrm{Tr}(d\mathbf{X}) \tag{18}$$

Combining the identities 14, 15, 18,above, we can obtain the second order derivatives of the log-likelihood in a similar way:

$$\begin{aligned}
\frac{\partial \mathcal{L}}{\partial \sigma_e^4} &= -(\widetilde{\mathbf{y}} - \widetilde{\mathbf{W}}\boldsymbol{\beta} - \widetilde{\mathbf{z}}\mu)^\top \widetilde{\mathbf{P}}\widetilde{\mathbf{P}}\widetilde{\mathbf{P}}(\widetilde{\mathbf{y}} - \widetilde{\mathbf{W}}\boldsymbol{\beta} - \widetilde{\mathbf{z}}\mu) + \frac{1}{2}\mathrm{Tr}(\widetilde{\mathbf{P}}\widetilde{\mathbf{P}}) \\
\frac{\partial \mathcal{L}}{\partial \sigma_e^2\sigma_1^2} &= -(\widetilde{\mathbf{y}} - \widetilde{\mathbf{W}}\boldsymbol{\beta} - \widetilde{\mathbf{z}}\mu)^\top \widetilde{\mathbf{P}}\widetilde{\mathbf{P}}\mathbf{S}_1\widetilde{\mathbf{P}}(\widetilde{\mathbf{y}} - \widetilde{\mathbf{W}}\boldsymbol{\beta} - \widetilde{\mathbf{z}}\mu) + \frac{1}{2}\mathrm{Tr}(\widetilde{\mathbf{P}}\widetilde{\mathbf{P}}\mathbf{S}_1) \\
\frac{\partial \mathcal{L}}{\partial \sigma_e^2\sigma_2^2} &= -(\widetilde{\mathbf{y}} - \widetilde{\mathbf{W}}\boldsymbol{\beta} - \widetilde{\mathbf{z}}\mu)^\top \widetilde{\mathbf{P}}\widetilde{\mathbf{P}}\mathbf{S}_2\widetilde{\mathbf{P}}(\widetilde{\mathbf{y}} - \widetilde{\mathbf{W}}_1\boldsymbol{\beta} - \widetilde{\mathbf{z}}\mu) + \frac{1}{2}\mathrm{Tr}(\widetilde{\mathbf{P}}\widetilde{\mathbf{P}}\mathbf{S}_2) \\
\frac{\partial \mathcal{L}}{\partial \sigma_1^4} &= -(\widetilde{\mathbf{y}} - \widetilde{\mathbf{W}}\boldsymbol{\beta} - \widetilde{\mathbf{z}}\mu)^\top \widetilde{\mathbf{P}}\mathbf{S}_1\widetilde{\mathbf{P}}\mathbf{S}_1\widetilde{\mathbf{P}}(\widetilde{\mathbf{y}} - \widetilde{\mathbf{W}}\boldsymbol{\beta} - \widetilde{\mathbf{z}}\mu) + \frac{1}{2}\mathrm{Tr}(\widetilde{\mathbf{P}}\mathbf{S}_1\widetilde{\mathbf{P}}\mathbf{S}_1) \\
\frac{\partial \mathcal{L}}{\partial \sigma_2^4} &= -(\widetilde{\mathbf{y}} - \widetilde{\mathbf{W}}\boldsymbol{\beta} - \widetilde{\mathbf{z}}\mu)^\top \widetilde{\mathbf{P}}\mathbf{S}_2\widetilde{\mathbf{P}}\mathbf{S}_2\widetilde{\mathbf{P}}(\widetilde{\mathbf{y}} - \widetilde{\mathbf{W}}\boldsymbol{\beta} - \widetilde{\mathbf{z}}\mu) + \frac{1}{2}\mathrm{Tr}(\widetilde{\mathbf{P}}\mathbf{S}_2\widetilde{\mathbf{P}}\mathbf{S}_2) \\
\frac{\partial \mathcal{L}}{\partial \sigma_1^2\sigma_2^2} &= -(\widetilde{\mathbf{y}} - \widetilde{\mathbf{W}}\boldsymbol{\beta} - \widetilde{\mathbf{z}}\mu)^\top \widetilde{\mathbf{P}}\mathbf{S}_1\widetilde{\mathbf{P}}\mathbf{S}_2\widetilde{\mathbf{P}}(\widetilde{\mathbf{y}} - \widetilde{\mathbf{W}}\boldsymbol{\beta} - \widetilde{\mathbf{z}}\mu) + \frac{1}{2}\mathrm{Tr}(\widetilde{\mathbf{P}}\mathbf{S}_1\widetilde{\mathbf{P}}\mathbf{S}_2)
\end{aligned} \tag{19}$$

Referring to the following expectation of quadratic form and assuming $\mathbf{A}$ is symmetric, $\mathbf{c} = \mathbb{E}[\mathbf{x}]$ and $\boldsymbol{\Sigma} = \mathrm{Var}[\mathbf{x}]$, then:

$$\mathbb{E}\left[\mathbf{x}^T\mathbf{A}\mathbf{x}\right] = \mathrm{Tr}(\mathbf{A}\boldsymbol{\Sigma}) + \mathbf{c}^T\mathbf{A}\mathbf{c} \tag{20}$$

Then we can obtain the expectation of the second order derivatives

$$
\begin{aligned}
\mathbb{E}\left[\frac{\partial L}{\partial \sigma_e^4}\right] &= \mathbb{E}\left[-(\widetilde{\mathbf{y}} - \widetilde{\mathbf{W}}\boldsymbol{\beta} - \widetilde{\mathbf{z}}\mu)^\top \widetilde{\mathbf{P}}\widetilde{\mathbf{P}}\widetilde{\mathbf{P}}(\widetilde{\mathbf{y}} - \widetilde{\mathbf{W}}\boldsymbol{\beta} - \widetilde{\mathbf{z}}\mu) + \frac{1}{2}\mathrm{Tr}(\widetilde{\mathbf{P}}\widetilde{\mathbf{P}})\right] \\
&= -\frac{1}{2}\mathrm{Tr}(\widetilde{\mathbf{P}}\widetilde{\mathbf{P}}) \\
\mathbb{E}\left[\frac{\partial L}{\partial \sigma_1^4}\right] &= \mathbb{E}\left[-(\widetilde{\mathbf{y}} - \widetilde{\mathbf{W}}\boldsymbol{\beta} - \widetilde{\mathbf{z}}\mu)^\top \widetilde{\mathbf{P}}\mathbf{S}_1\widetilde{\mathbf{P}}\mathbf{S}_1\widetilde{\mathbf{P}}(\widetilde{\mathbf{y}} - \widetilde{\mathbf{W}}\boldsymbol{\beta} - \widetilde{\mathbf{z}}\mu) + \frac{1}{2}\mathrm{Tr}(\widetilde{\mathbf{P}}\mathbf{S}_1\widetilde{\mathbf{P}}\mathbf{S}_1)\right] \\
&= -\frac{1}{2}\mathrm{Tr}(\widetilde{\mathbf{P}}\mathbf{S}_1\widetilde{\mathbf{P}}\mathbf{S}_1) \\
\mathbb{E}\left[\frac{\partial L}{\partial \sigma_2^4}\right] &= \mathbb{E}\left[-(\widetilde{\mathbf{y}} - \widetilde{\mathbf{W}}\boldsymbol{\beta} - \widetilde{\mathbf{z}}\mu)^\top \widetilde{\mathbf{P}}\mathbf{s}_2\widetilde{\mathbf{P}}\mathbf{S}_2\widetilde{\mathbf{P}}(\widetilde{\mathbf{y}} - \widetilde{\mathbf{W}}\boldsymbol{\beta} - \widetilde{\mathbf{z}}\mu) + \frac{1}{2}\mathrm{Tr}(\widetilde{\mathbf{P}}\mathbf{S}_2\widetilde{\mathbf{P}}\mathbf{S}_2)\right] \\
&= -\frac{1}{2}\mathrm{Tr}(\widetilde{\mathbf{P}}\mathbf{S}_2\widetilde{\mathbf{P}}\mathbf{S}_2)
\end{aligned}
\tag{21}
$$

and

$$
\begin{aligned}
\mathbb{E}\left[\frac{\partial L}{\partial \sigma_e^2\sigma_1^2}\right] &= \mathbb{E}\left[-(\widetilde{\mathbf{y}} - \widetilde{\mathbf{W}}\boldsymbol{\beta} - \widetilde{\mathbf{z}}\mu)^\top \widetilde{\mathbf{P}}\widetilde{\mathbf{P}}\mathbf{S}_1\widetilde{\mathbf{P}}(\widetilde{\mathbf{y}} - \widetilde{\mathbf{W}}\boldsymbol{\beta} - \widetilde{\mathbf{z}}\mu) + \frac{1}{2}\mathrm{Tr}(\widetilde{\mathbf{P}}\widetilde{\mathbf{P}}\mathbf{S}_1)\right] \\
&= -\frac{1}{2}\mathrm{Tr}(\widetilde{\mathbf{P}}\widetilde{\mathbf{P}}\mathbf{S}_1) \\
\mathbb{E}\left[\frac{\partial L}{\partial \sigma_e^2\sigma_2^2}\right] &= \mathbb{E}\left[-(\widetilde{\mathbf{y}} - \widetilde{\mathbf{W}}\boldsymbol{\beta} - \widetilde{\mathbf{z}}\mu)^\top \widetilde{\mathbf{P}}\widetilde{\mathbf{P}}\mathbf{S}_2\widetilde{\mathbf{P}}(\widetilde{\mathbf{y}} - \widetilde{\mathbf{W}}_1\boldsymbol{\beta} - \widetilde{\mathbf{z}}\mu) + \frac{1}{2}\mathrm{Tr}(\widetilde{\mathbf{P}}\widetilde{\mathbf{P}}\mathbf{S}_2)\right] \\
&= -\frac{1}{2}\mathrm{Tr}(\widetilde{\mathbf{P}}\widetilde{\mathbf{P}}\mathbf{S}_2) \\
\mathbb{E}\left[\frac{\partial L}{\partial \sigma_1^2\sigma_2^2}\right] &= \mathbb{E}\left[-(\widetilde{\mathbf{y}} - \widetilde{\mathbf{W}}\boldsymbol{\beta} - \widetilde{\mathbf{z}}\mu)^\top \widetilde{\mathbf{P}}\mathbf{S}_1\widetilde{\mathbf{P}}\mathbf{S}_2\widetilde{\mathbf{P}}(\widetilde{\mathbf{y}} - \widetilde{\mathbf{W}}\boldsymbol{\beta} - \widetilde{\mathbf{z}}\mu) + \frac{1}{2}\mathrm{Tr}(\widetilde{\mathbf{P}}\mathbf{S}_1\widetilde{\mathbf{P}}\mathbf{S}_2)\right] \\
&= -\frac{1}{2}\mathrm{Tr}(\widetilde{\mathbf{P}}\mathbf{S}_1\widetilde{\mathbf{P}}\mathbf{S}_2)
\end{aligned}
\tag{22}
$$

By averaging the second order derivatives and their expectations, we can obtain the AI matrix

$$
\begin{aligned}
\mathrm{AI}(\sigma_e^2, \sigma_e^2) &= -(\widetilde{\mathbf{y}} - \widetilde{\mathbf{W}}\boldsymbol{\beta} - \widetilde{\mathbf{z}}\mu)^\top \widetilde{\mathbf{P}}\widetilde{\mathbf{P}}\widetilde{\mathbf{P}}(\widetilde{\mathbf{y}} - \widetilde{\mathbf{W}}\boldsymbol{\beta} - \widetilde{\mathbf{z}}\mu) \\
\mathrm{AI}(\sigma_1^2, \sigma_1^2) &= -(\widetilde{\mathbf{y}} - \widetilde{\mathbf{W}}\boldsymbol{\beta} - \widetilde{\mathbf{z}}\mu)^\top \widetilde{\mathbf{P}}\mathbf{S}_1\widetilde{\mathbf{P}}\mathbf{S}_1\widetilde{\mathbf{P}}(\widetilde{\mathbf{y}} - \widetilde{\mathbf{W}}\boldsymbol{\beta} - \widetilde{\mathbf{z}}\mu) \\
\mathrm{AI}(\sigma_2^2, \sigma_2^2) &= -(\widetilde{\mathbf{y}} - \widetilde{\mathbf{W}}\boldsymbol{\beta} - \widetilde{\mathbf{z}}\mu)^\top \widetilde{\mathbf{P}}\mathbf{S}_2\widetilde{\mathbf{P}}\mathbf{S}_2\widetilde{\mathbf{P}}(\widetilde{\mathbf{y}} - \widetilde{\mathbf{W}}\boldsymbol{\beta} - \widetilde{\mathbf{z}}\mu) \\
\mathrm{AI}(\sigma_e^2, \sigma_1^2) &= -(\widetilde{\mathbf{y}} - \widetilde{\mathbf{W}}\boldsymbol{\beta} - \widetilde{\mathbf{z}}\mu)^\top \widetilde{\mathbf{P}}\widetilde{\mathbf{P}}\mathbf{S}_1\widetilde{\mathbf{P}}(\widetilde{\mathbf{y}} - \widetilde{\mathbf{W}}\boldsymbol{\beta} - \widetilde{\mathbf{z}}\mu) \\
\mathrm{AI}(\sigma_e^2, \sigma_2^2) &= -(\widetilde{\mathbf{y}} - \widetilde{\mathbf{W}}\boldsymbol{\beta} - \widetilde{\mathbf{z}}\mu)^\top \widetilde{\mathbf{P}}\widetilde{\mathbf{P}}\mathbf{S}_2\widetilde{\mathbf{P}}(\widetilde{\mathbf{y}} - \widetilde{\mathbf{W}}\boldsymbol{\beta} - \widetilde{\mathbf{z}}\mu) \\
\mathrm{AI}(\sigma_1^2, \sigma_2^2) &= -(\widetilde{\mathbf{y}} - \widetilde{\mathbf{W}}\boldsymbol{\beta} - \widetilde{\mathbf{z}}\mu)^\top \widetilde{\mathbf{P}}\mathbf{S}_1\widetilde{\mathbf{P}}\mathbf{S}_2\widetilde{\mathbf{P}}(\widetilde{\mathbf{y}} - \widetilde{\mathbf{W}}\boldsymbol{\beta} - \widetilde{\mathbf{z}}\mu)
\end{aligned}
\tag{23}
$$

We use the Newton-Raphson algorithm to update parameters $\boldsymbol{\theta}_1 = \{\sigma_1^2, \sigma_2^2, \sigma_e^2\}$

$$
\boldsymbol{\theta}_1^{t+1} = \boldsymbol{\theta}_1^t - \mathrm{AI}^{-1}\frac{\partial \mathcal{L}}{\partial \boldsymbol{\theta}_1} \tag{24}
$$

For parameters $\boldsymbol{\beta}$ and $\mu$, we derive the closed forms of these parameters by setting the first derivatives of the log-likelihood with respect to $\boldsymbol{\beta}$ and $\mu$ equal to 0.

$$
\begin{aligned}
\frac{\partial \mathcal{L}}{\partial \boldsymbol{\beta}} &= -\widetilde{\mathbf{W}}^\top \mathbf{P}(\widetilde{\mathbf{y}} - \widetilde{\mathbf{W}}\boldsymbol{\beta} - \widetilde{\mathbf{z}}\mu) = \mathbf{0} \\
\frac{\partial \mathcal{L}}{\partial \mu} &= -\widetilde{\mathbf{W}}^\top \mathbf{P}(\widetilde{\mathbf{y}} - \widetilde{\mathbf{W}}\boldsymbol{\beta} - \widetilde{\mathbf{z}}\mu) = \mathbf{0}
\end{aligned} \tag{25}
$$

The closed forms of $\boldsymbol{\beta}$ and $\mu$ are

$$
\begin{aligned}
\boldsymbol{\beta} &= (\widetilde{\mathbf{W}}^\top \widetilde{\mathbf{P}} \widetilde{\mathbf{W}})^{-1} \widetilde{\mathbf{W}}^\top \widetilde{\mathbf{P}}(\widetilde{\mathbf{y}} - \widetilde{\mathbf{z}}\mu) \\
\mu &= (\widetilde{\mathbf{z}}^\top \widetilde{\mathbf{P}} \widetilde{\mathbf{z}})^{-1} \widetilde{\mathbf{z}}^\top \widetilde{\mathbf{P}}(\widetilde{\mathbf{y}} - \widetilde{\mathbf{W}}\boldsymbol{\beta})
\end{aligned} \tag{26}
$$

Finally, the whole algorithm is summarized as Algorithm 1.

#### 1.6 Algorithm of spatialDEG

**Algorithm 1**: AI algorithm

Initialize $\boldsymbol{\theta}_1 = \{\sigma_1^2, \sigma_2^2, \sigma_e^2\}, \boldsymbol{\beta}, \mu$, given a predefined sequence of $l_1, \cdots, l_{k_1}$ and $\phi_1, \cdots, \phi_{k_2}$ for the Gaussian and Cosine kernels, respectively.

Precalculate the Eigen decomposition of $\mathbf{K}_1$ and $\mathbf{K}_2$ with the length scale $l_1, \cdots, l_{k_1}$, and periodicity parameter $\phi_1, \cdots, \phi_{k_2}$.

Set $t = 1$, $\phi = l_1$.

**repeat**

  update $\widetilde{\mathbf{y}} = \mathbf{U}^\top \mathbf{y}$, $\widetilde{\mathbf{W}} = \mathbf{U}^\top \mathbf{W}$, $\widetilde{\mathbf{z}} = \mathbf{U}^\top \mathbf{z}$

  $\boldsymbol{\beta} = (\widetilde{\mathbf{W}}^\top \widetilde{\mathbf{P}} \widetilde{\mathbf{W}})^{-1} \widetilde{\mathbf{W}}^\top \widetilde{\mathbf{P}}(\widetilde{\mathbf{y}} - \widetilde{\mathbf{z}}\mu)$

  $\mu = (\widetilde{\mathbf{z}}^\top \widetilde{\mathbf{P}} \widetilde{\mathbf{z}})^{-1} \widetilde{\mathbf{z}}^\top \widetilde{\mathbf{P}}(\widetilde{\mathbf{y}} - \widetilde{\mathbf{W}}\boldsymbol{\beta})$

  $\boldsymbol{\theta}_1^{t+1} = \boldsymbol{\theta}_1^t - \mathrm{AI}^{-1} \frac{\partial \mathcal{L}(\boldsymbol{\theta}_1)}{\partial \boldsymbol{\theta}_1}$, where AI is defined as equation (23)

  $\widehat{\phi} = \arg\max_{\phi \in \{l_1, \cdots, l_{k_1}, \phi_1, \cdots, \phi_{k_2}\}} \mathcal{L}(\phi)$

  $t = t + 1$

**until** *the change of $\mathcal{L}(\boldsymbol{\theta})$ is smaller than a threshold*;

**return** $\boldsymbol{\theta}$ *and* $\phi$

### 2 Simulation studies

#### 2.1 Simulation setting and data generation

In scenario (a), for the first dataset, the spatial locations of the spots were taken from the real data sample HFD1 or WT2. The number of spots for samples HFD1 and WT2 were $n_1 = 1328$ and $n_1 = 3613$, respectively, and the number of genes was set to 1000. The simulated gene expression was the summation of two components: the spatial component and the non-spatial component. The spatial component was generated from Gaussian distribution with a mean of zero and a variance-covariance matrix Gaussian kernel or Cosine kernel. We chose the 0.1st, 0.3rd, 0.5th, 0.7th and 0.9th quantiles of all the pairwise Euclidean distances of all spots as the length scale of the Gaussian kernels or the periodicity parameters of the Cosine kernels. These were 10.2, 19.7, 27.86, 37.2, and 53.14 for HFD1 and 16.55, 31.33, 44, 57.98 and 79.21 for WT2, respectively. The magnitude of the kernel was set to $\sigma_1^2 = 1$. The non-spatial component was generated from the Gaussian distribution with a mean of zero and a variance-covariance matrix identity matrix. The magnitude of the identity matrix was set to $\sigma_e^2 = 1$. We referred to the ratio of the variance of the spatial component relative to the summation of the variance of the spatial component and the variance of the non-spatial component as the heritability, which was denoted as $h^2$, which was controlled at 0.1, 0.2, and 0.3. Then the second dataset was generated to be identical to the first dataset. The intercept was set to 5.

In scenario (b), both datasets in each scenario were generated to be similar to the null hypothesis; the only difference was that the spatial component was generated from a Gaussian distribution with a mean $Diff$ and variance-covariance matrix Gaussian kernel or Cosine kernel in the second dataset. The $Diff$ was set to 0, 0.05, 0.1, 0.15, and 0.2 for HFD1 and 0, 0.25, 0.5, 0.75, and 1 for WT2.

In this simulation study, spatialDEG was used with both the true kernel and estimated kernel separately in all settings to further explore the effects of the kernel on the type-1 error rate and power. The number of the length scale or periodicity were set to $k_1 = k_2 = 40$ for HFD1 and $k_1 = k_2 = 60$ for WT2.

#### 2.2 Summary of simulation settings

In order to demonstrate the simulation settings clearly, we summarize the simulation settings for scenario (a) and scenario (b) in the following tables:

| Type of kernel | Spatial location | $h^2 = 0.1$ | $h^2 = 0.2$ | $h^2 = 0.3$ |
|---|---|---|---|---|
| Gaussian | HFD1 | Figure 1 | Figure 2 | Figure 3 |
| Gaussian | WT2 | Figure 4 | Figure 5 | Figure 6 |
| Cosine | HFD1 | Figure 7 | Figure 8 | Figure 9 |
| Cosine | WT2 | Figure 10 | Figure 11 | Figure 12 |

Table 1: Simulation settings for scenario (a), in which spatialDEG is used with true kernel.

| Type of kernel | Spatial location | $h^2 = 0.1$ | $h^2 = 0.2$ | $h^2 = 0.3$ |
|---|---|---|---|---|
| Gaussian | HFD1 | Figure 13 | Figure 14 | Figure 15 |
| Gaussian | WT2 | Figure 16 | Figure 17 | Figure 18 |
| Cosine | HFD1 | Figure 19 | Figure 20 | Figure 21 |
| Cosine | WT2 | Figure 22 | Figure 23 | Figure 24 |

Table 2: Simulation settings for scenario (a), in which spatialDEG is used with the estimated kernel.

| True or estimated kernel | Type of kernel | HFD1 | WT2 |
|---|---|---|---|
| True | Gaussian | Figure 25 | Figure 26 |
| True | Cosine | Figure 27 | Figure 28 |
| Estimated | Gaussian | Figure 29 | Figure 30 |
| Estimated | Cosine | Figure 31 | Figure 32 |

Table 3: Simulation settings for the scenario (b).

#### 2.3 Type 1 error

##### 2.3.1 The ground truth is the Gaussian kernel, and SpatialDEG is used with the true Gaussian kernel

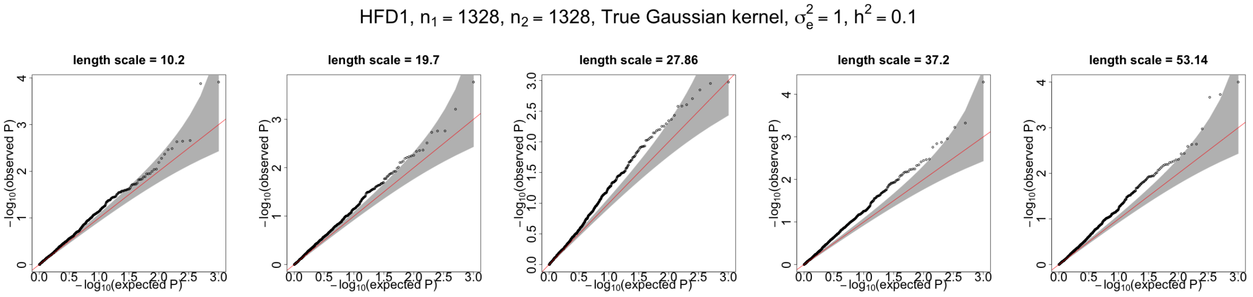


Figure 1: qq-plots for p-values generated by spatialDEG with the true Gaussian kernel. Spatial location is based on HFD1 data. The ground truth is the Gaussian kernel. Heritability $h^2 = 0.1$.

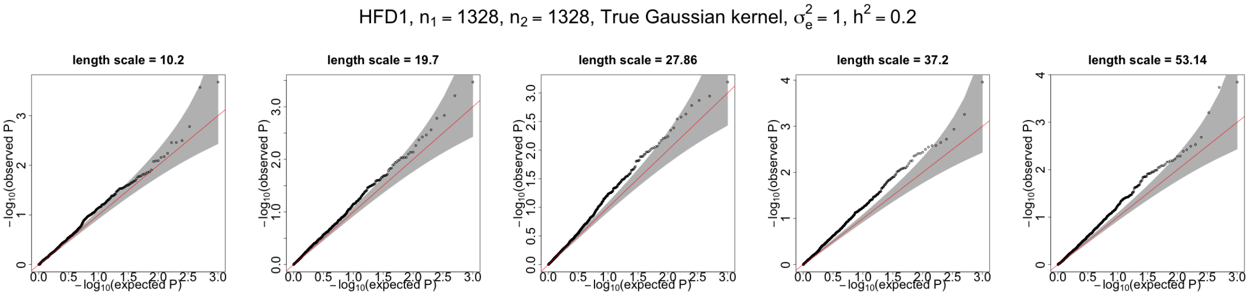


Figure 2: qq-plots for p-values generated by spatialDEG with the true Gaussian kernel. Spatial location is based on HFD1 data. The ground truth is the Gaussian kernel. Heritability $h^2 = 0.2$.

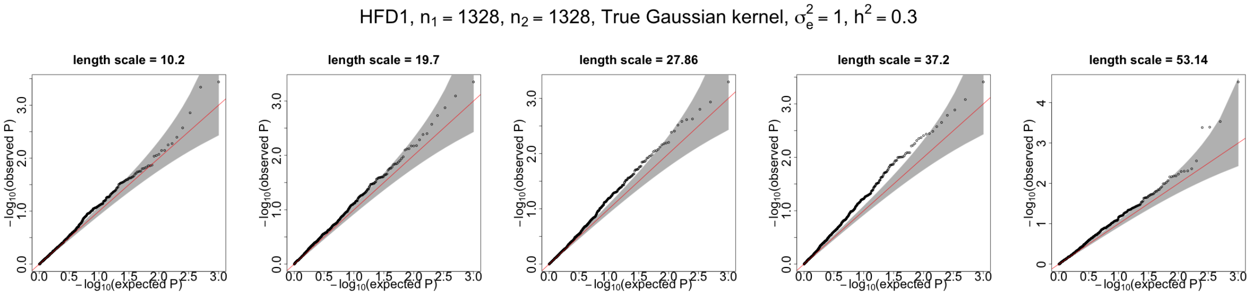


Figure 3: qq-plots for p-values generated by spatialDEG with the true Gaussian kernel. Spatial location is based on HFD1 data. The ground truth is the Gaussian kernel. Heritability $h^2 = 0.3$.

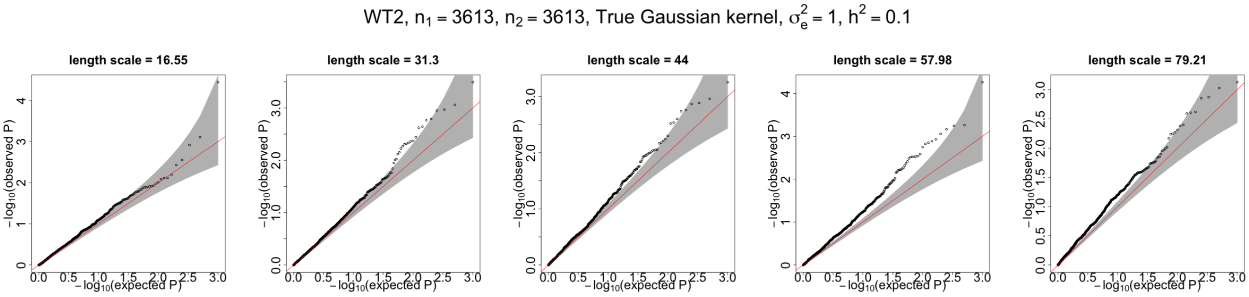


Figure 4: qq-plots for p-values generated by spatialDEG with the true Gaussian kernel. Spatial location is based on WT2 data. The ground truth is the Gaussian kernel. Heritability $h^2 = 0.1$.

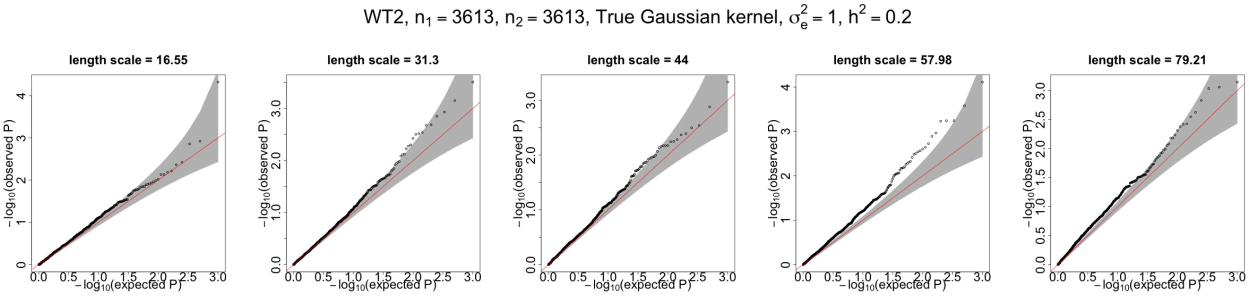


Figure 5: qq-plots for p-values generated by spatialDEG with the true Gaussian kernel. Spatial location is based on WT2 data. The ground truth is the Gaussian kernel. Heritability $h^2 = 0.2$.

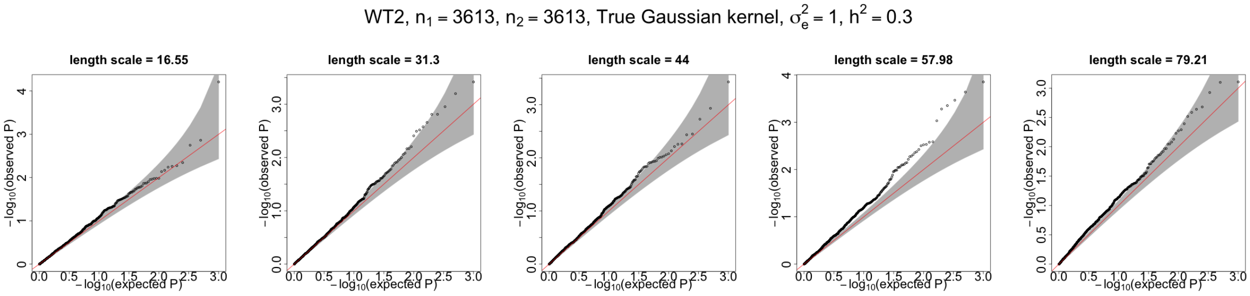


Figure 6: qq-plots for p-values generated by spatialDEG with the true Gaussian kernel. Spatial location is based on WT2 data. The ground truth is the Gaussian kernel. Heritability $h^2 = 0.3$.

##### 2.3.2 The ground truth is Cosine kernel. SpatialDEG is used with the true Cosine kernel.

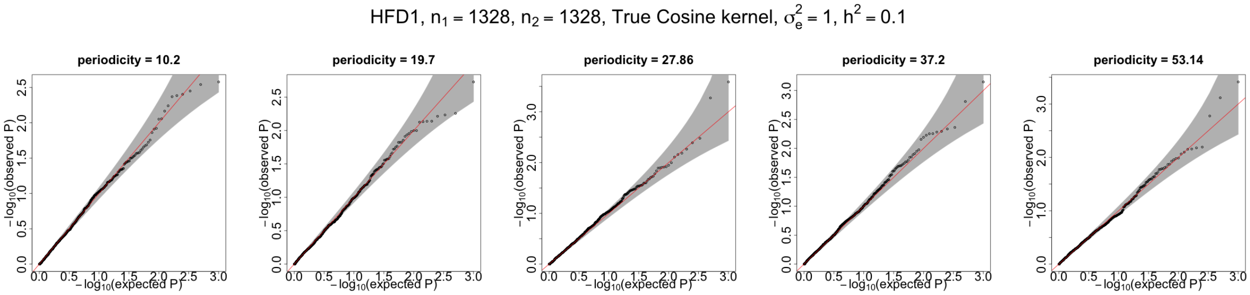


Figure 7: qq-plots for p-values generated by spatialDEG with the true Cosine kernel. Spatial location is based on HFD1 data. The ground truth is the Cosine kernel. Heritability $h^2 = 0.1$.

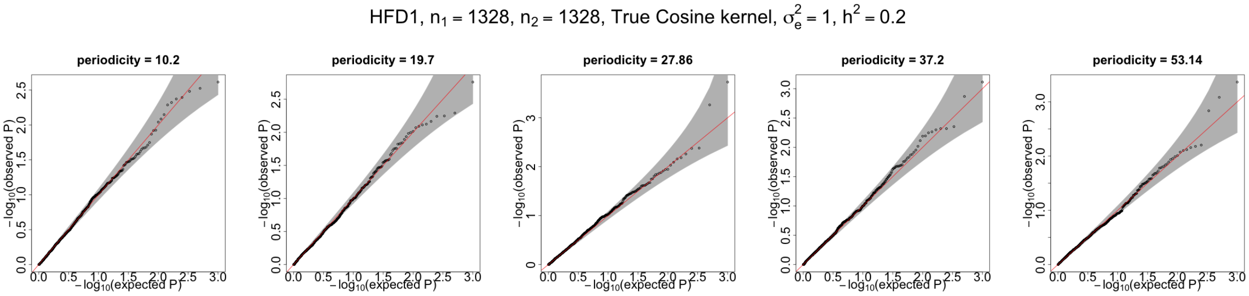


Figure 8: qq-plots for p-values generated by spatialDEG with the true Cosine kernel. Spatial location is based on HFD1 data. The ground truth is the Cosine kernel. Heritability $h^2 = 0.2$.

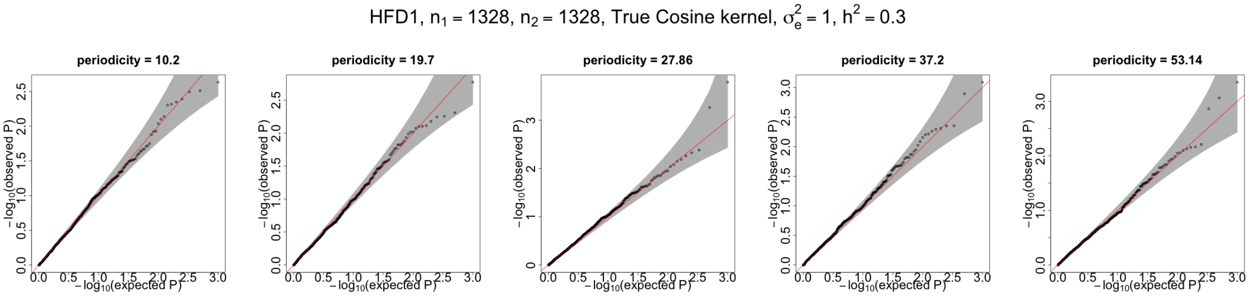


Figure 9: qq-plots for p-values generated by spatialDEG with the true Cosine kernel. Spatial location is based on HFD1 data. The ground truth is the Cosine kernel. Heritability $h^2 = 0.3$.

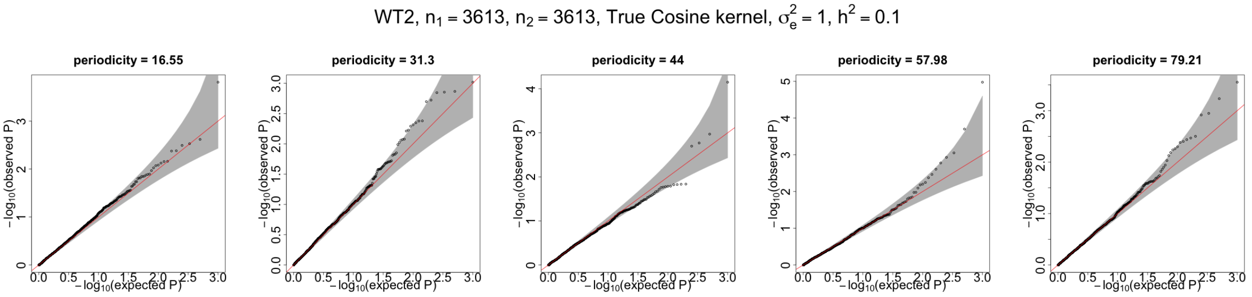


Figure 10: qq-plots for p-values generated by spatialDEG with the true Cosine kernel. Spatial location is based on WT2 data. The ground truth is the Cosine kernel. Heritability $h^2 = 0.1$.

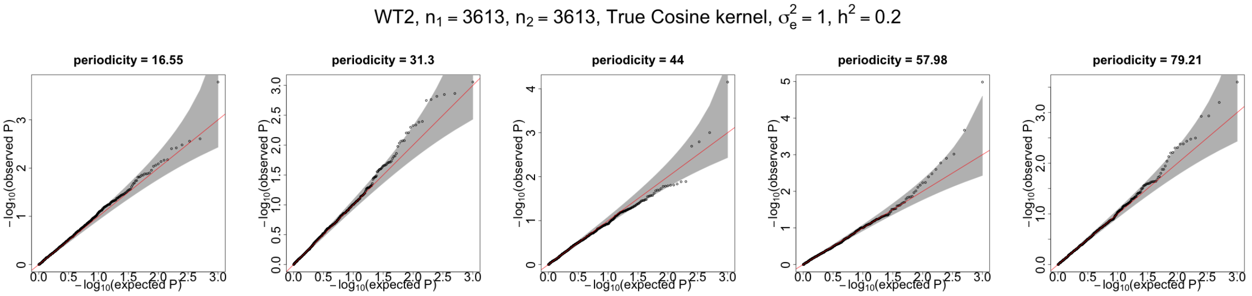


Figure 11: qq-plots for p-values generated by spatialDEG with the true Cosine kernel. Spatial location is based on WT2 data. The ground truth is the Cosine kernel. Heritability $h^2 = 0.2$.

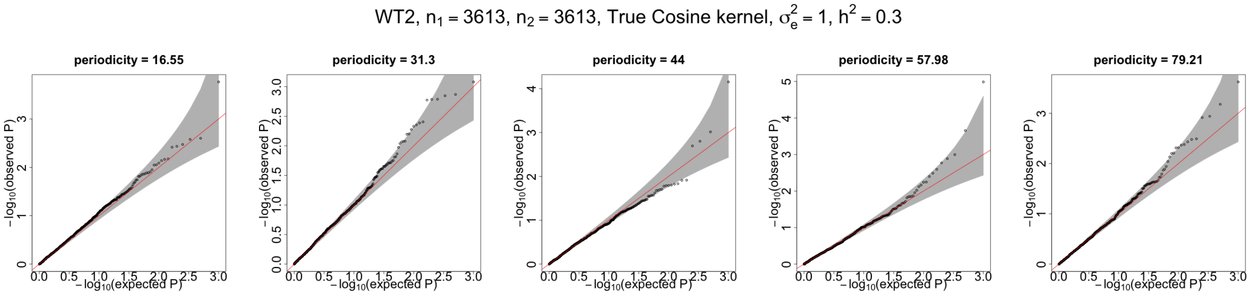


Figure 12: qq-plots for p-values generated by spatialDEG with the true Cosine kernel. Spatial location is based on WT2 data. The ground truth is the Cosine kernel. Heritability $h^2 = 0.3$.

##### 2.3.3 The ground truth is the Gaussian kernel. SpatialDEG is used with the estimated kernel.

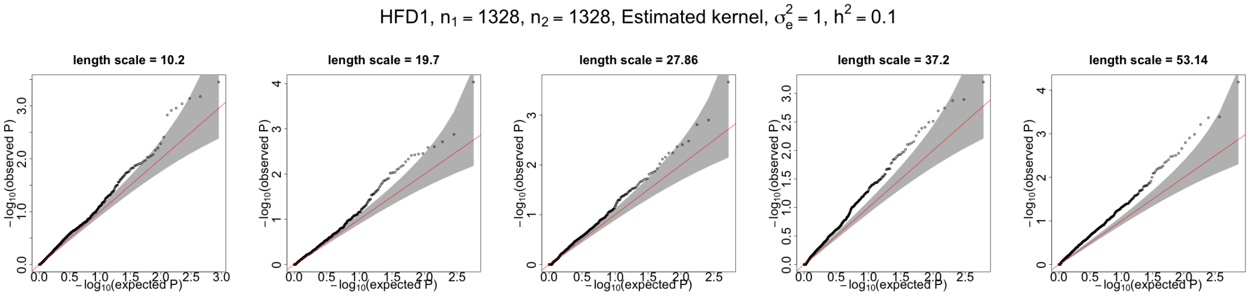


Figure 13: qq-plots for p-values generated by spatialDEG with the estimated kernel. Spatial location is based on HFD1 data. The ground truth is the Gaussian kernel. Heritability $h^2 = 0.1$.

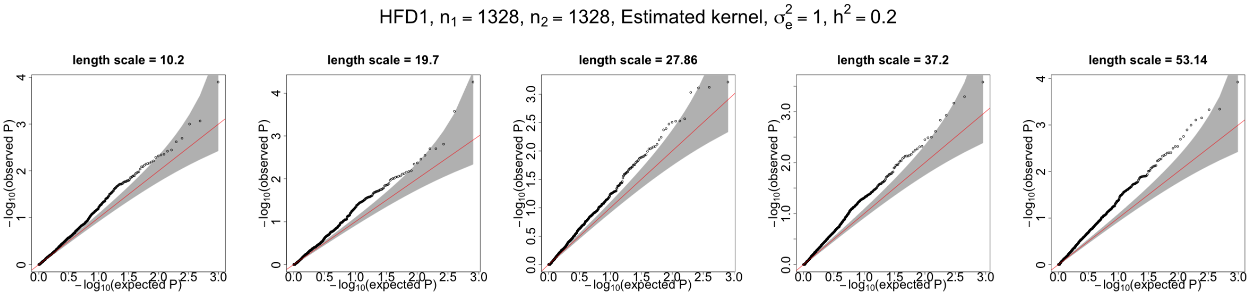


Figure 14: qq-plots for p-values generated by spatialDEG with the estimated kernel. Spatial location is based on HFD1 data. The ground truth is the Gaussian kernel. Heritability $h^2 = 0.2$.

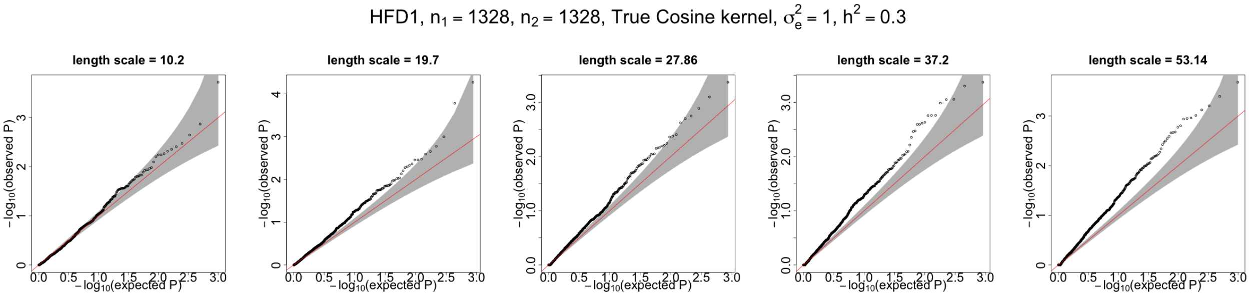


Figure 15: qq-plots for p-values generated by spatialDEG with the estimated kernel. Spatial location is based on HFD1 data. The ground truth is the Gaussian kernel. Heritability $h^2 = 0.3$.

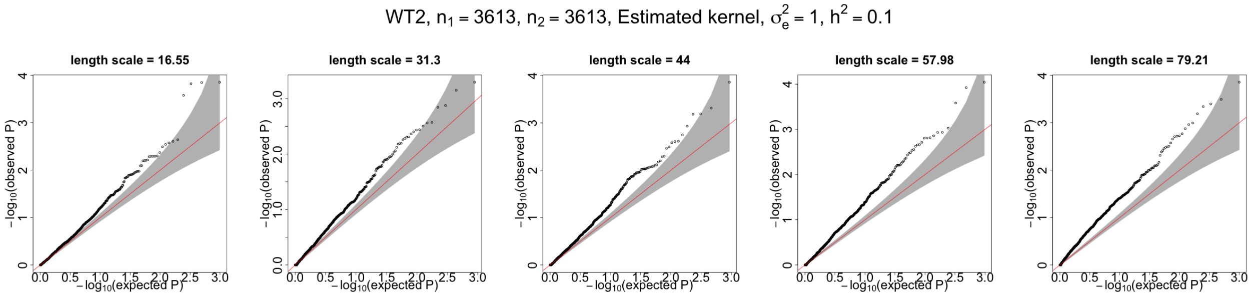


Figure 16: qq-plots for p-values generated by spatialDEG with the estimated kernel. Spatial location is based on WT2 data. The ground truth is the Gaussian kernel. Heritability $h^2 = 0.1$.

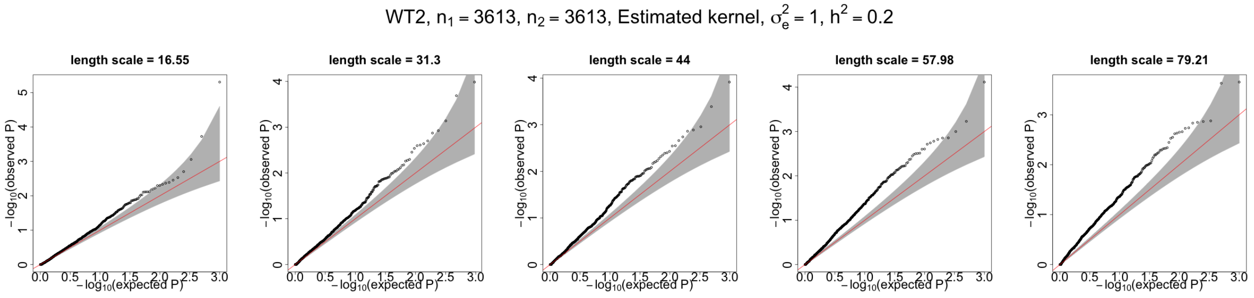


Figure 17: qq-plots for p-values generated by spatialDEG with the estimated kernel. Spatial location is based on WT2 data. The ground truth is the Gaussian kernel. Heritability $h^2 = 0.2$.

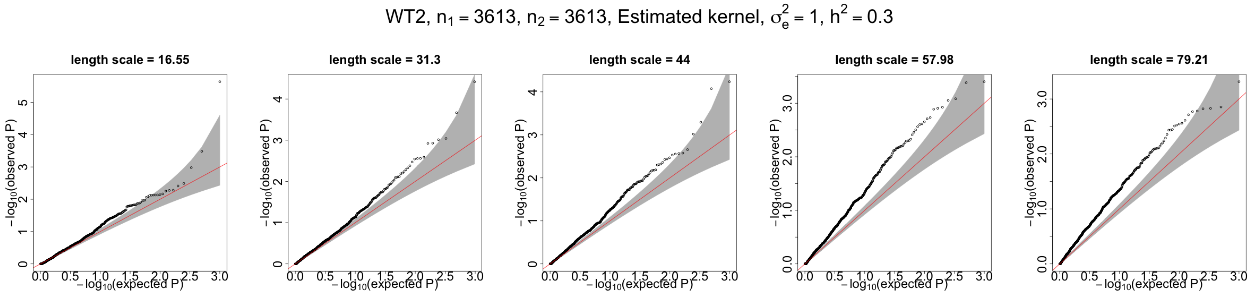


Figure 18: qq-plots for p-values generated by spatialDEG with the estimated kernel. Spatial location is based on WT2 data. The ground truth is the Gaussian kernel. Heritability $h^2 = 0.3$.

##### 2.3.4 The ground truth is the Cosine kernel. SpatialDEG is used with the estimated kernel.

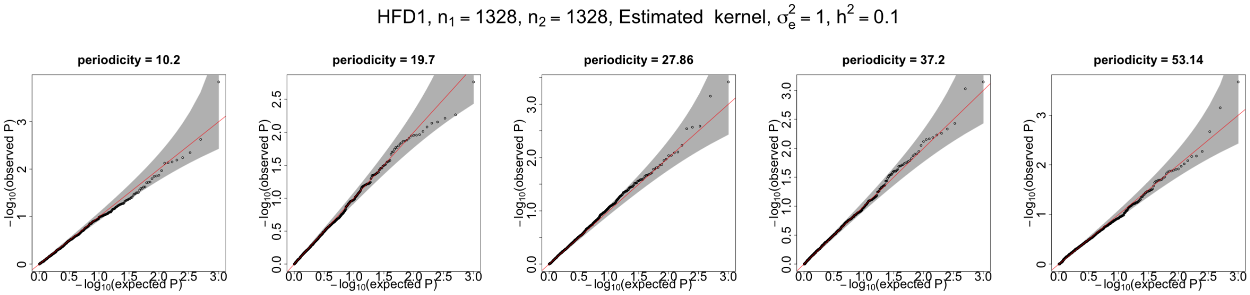


Figure 19: qq-plots for p-values generated by spatialDEG with the estimated kernel. Spatial location is based on HFD1 data. The ground truth is the Cosine kernel. Heritability $h^2 = 0.1$.

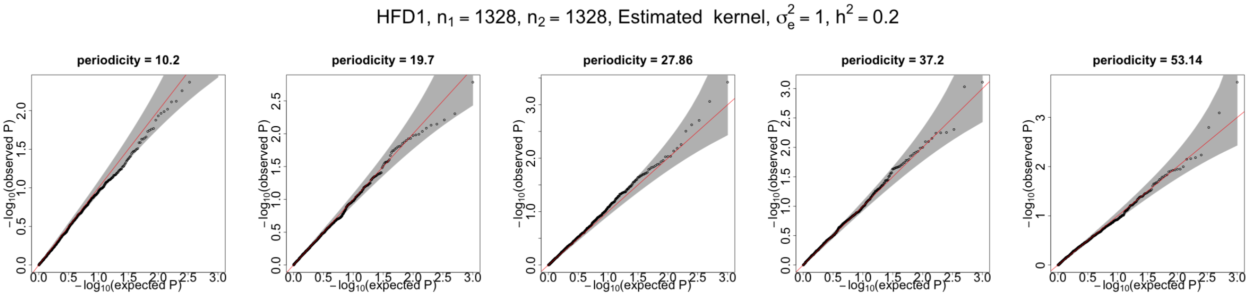


Figure 20: qq-plots for p-values generated by spatialDEG with the estimated kernel. Spatial location is based on HFD1 data. The ground truth is the Cosine kernel. Heritability $h^2 = 0.2$.

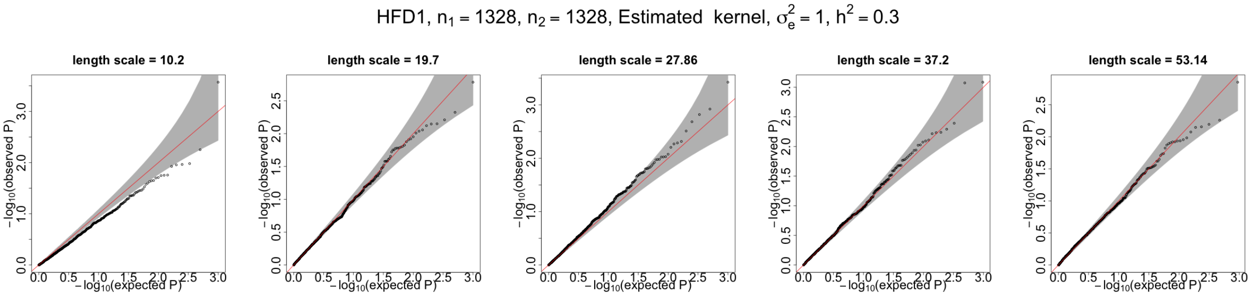


Figure 21: qq-plots for p-values generated by spatialDEG with the estimated kernel. Spatial location is based on HFD1 data. The ground truth is the Cosine kernel. Heritability $h^2 = 0.3$.

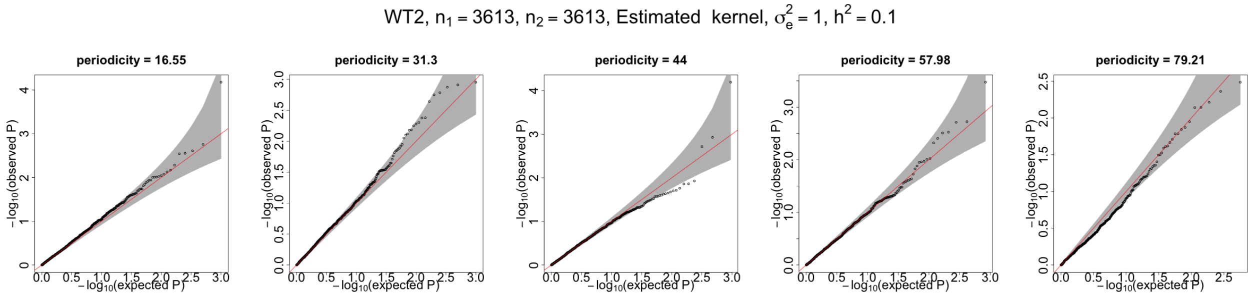


Figure 22: qq-plots for p-values generated by spatialDEG with the estimated kernel. Spatial location is based on WT2 data. The ground truth is the Cosine kernel. Heritability $h^2 = 0.1$.

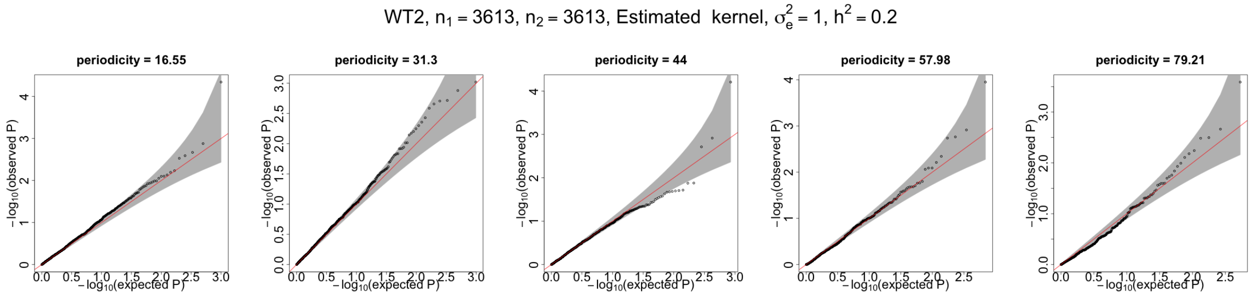


Figure 23: qq-plots for p-values generated by spatialDEG with the estimated kernel. Spatial location is based on WT2 data. The ground truth is the Cosine kernel. Heritability $h^2 = 0.2$.

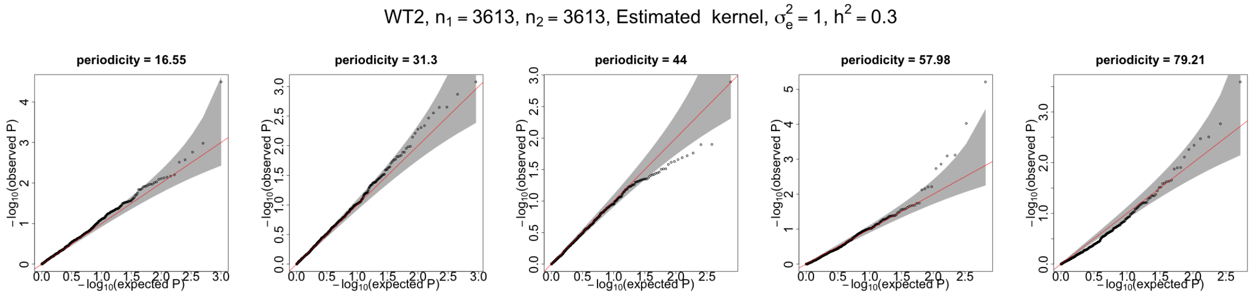


Figure 24: qq-plots for p-values generated by spatialDEG with the estimated kernel. Spatial location is based on WT2 data. The ground truth is the Cosine kernel. Heritability $h^2 = 0.3$.

#### 2.4 Power

##### 2.4.1 The ground truth is the Gaussian kernel. SpatialDEG is used with the true Gaussian kernel

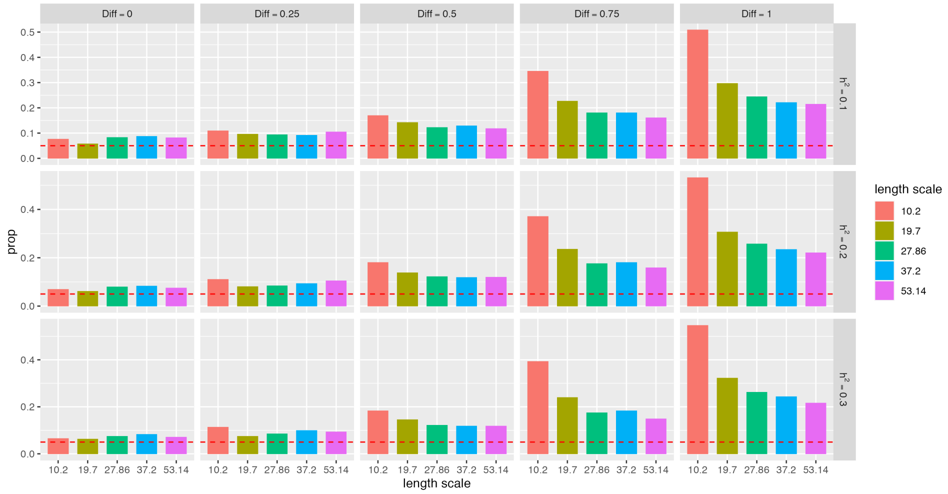


Figure 25: Bar plots for the power of spatialDEG with the true Gaussian kernel. The ground truth is the Gaussian kernel. The x axis provides the length scale, and the y axis provides the proportion (prop) of significant genes. Diff denotes the difference in the mean of the spatial component between two datasets. The spot location is based on HFD1. The number of spots $n_1 = n_2 = 1328$.

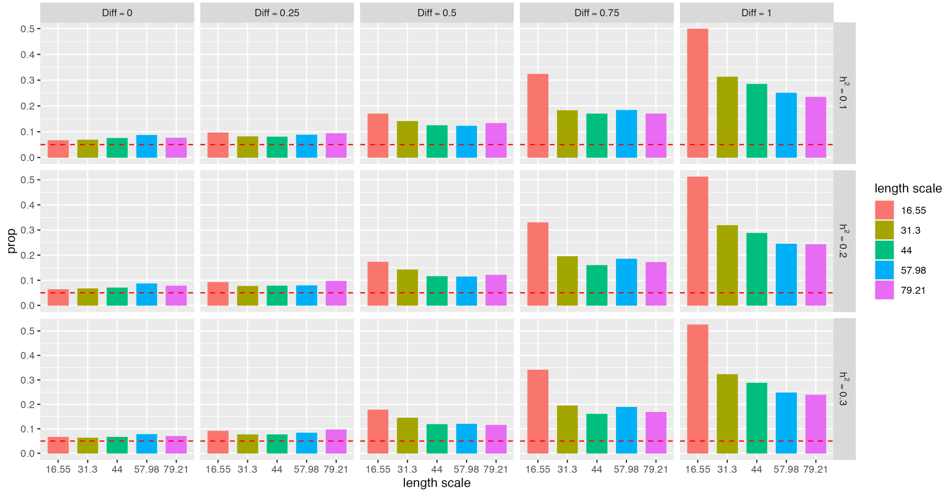


Figure 26: Bar plots for the power of spatialDEG with the true Gaussian kernel. The ground truth is the Gaussian kernel. The x axis provides the length scale, and the y axis provides the proportion (prop) of significant genes. Diff denotes the difference in the mean of the spatial component between two datasets. The spot location is based on WT2. The number of spots $n_1 = n_2 = 3613$.

##### 2.4.2 The ground truth is the Cosine kernel. SpatialDEG is used with the true Cosine kernel.

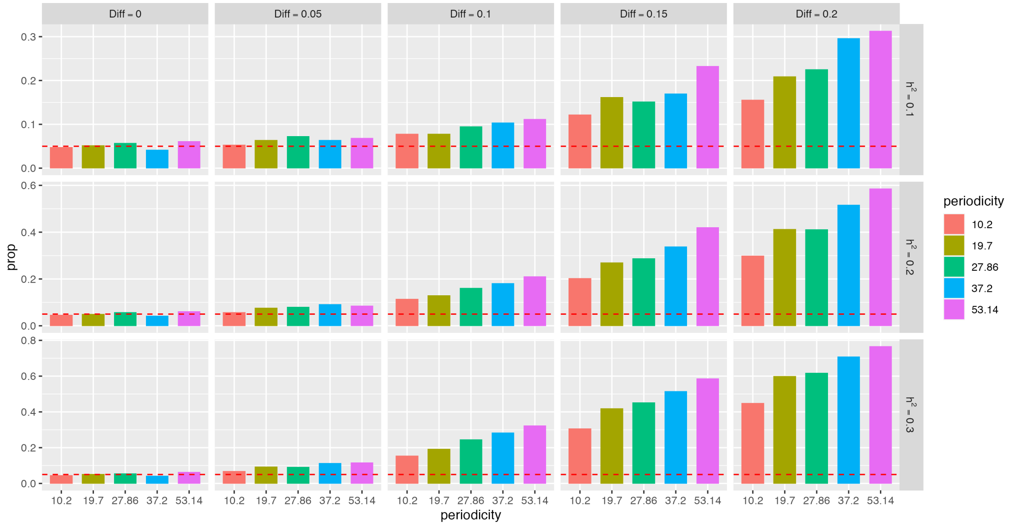


Figure 27: Bar plots for the power of spatialDEG with the true cosine kernel. The ground truth is the Cosine kernel. The x axis provides the periodicity parameters, and the y axis provides the proportion (prop) of significant genes. Diff denotes the difference in the mean of the spatial component between two datasets. The spot location is based on HFD1. The number of spots $n_1 = n_2 = 1328$.

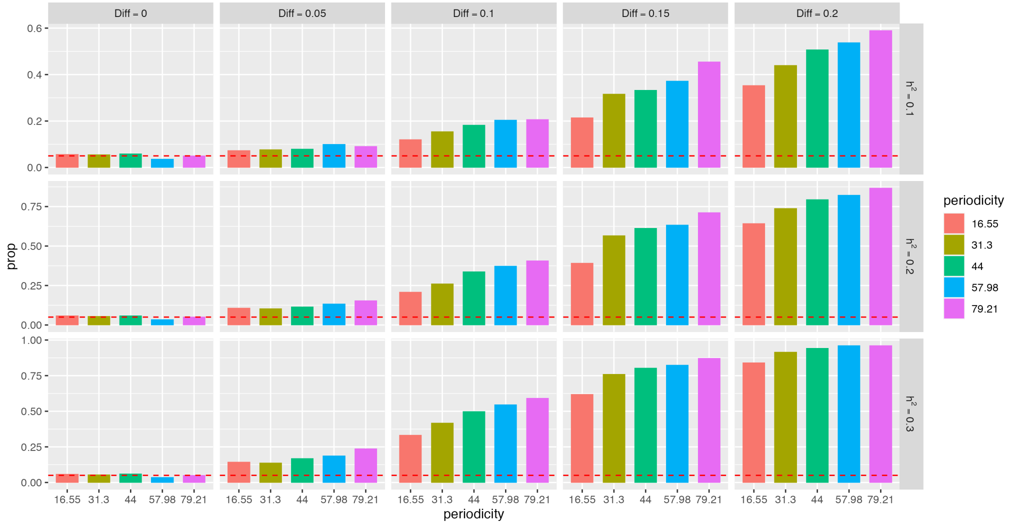


Figure 28: Bar plots for the power of spatialDEG with the true cosine kernel. The ground truth is the Cosine kernel. The x axis provides the periodicity parameters, and the y axis provides the proportion (prop) of significant genes. Diff denotes the difference in the mean of the spatial component between two datasets. The spot location is based on WT2. The number of spots is $n_1 = n_2 = 3613$.

##### 2.4.3 The ground truth is the Gaussian kernel. SpatialDEG is used with the estimated kernel.

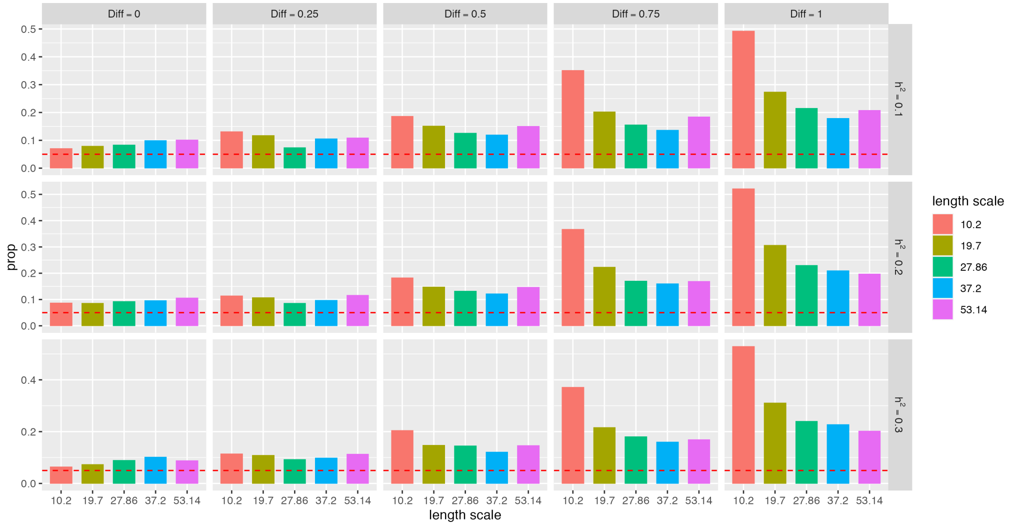


Figure 29: Bar plots for the power of spatialDEG with the estimated kernel. The ground truth is the Gaussian kernel. The x axis provides the length scale, and the y axis provides the proportion (prop) of significant genes. Diff denotes the difference in the means of the spatial components between the two datasets. The spot location is based on HFD1. The number of spots $n_1 = n_2 = 1328$.

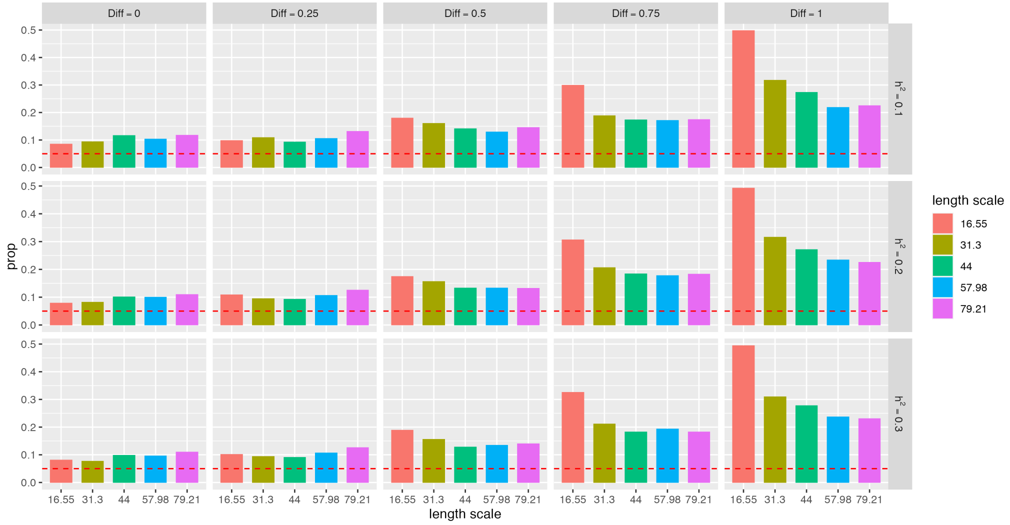


Figure 30: Bar plots for the power of spatialDEG with the estimated kernel. The ground truth is the Gaussian kernel. The x axis provides the length scale and the y axis provides the proportion (prop) of significant genes. Diff denotes the difference in the means of the spatial components between the two datasets. The spot location is based on WT2. The number of spots is $n_1 = n_2 = 3613$.

###### 2.4.4 The ground truth is the Cosine kernel. SpatialDEG is used with the estimated kernel.

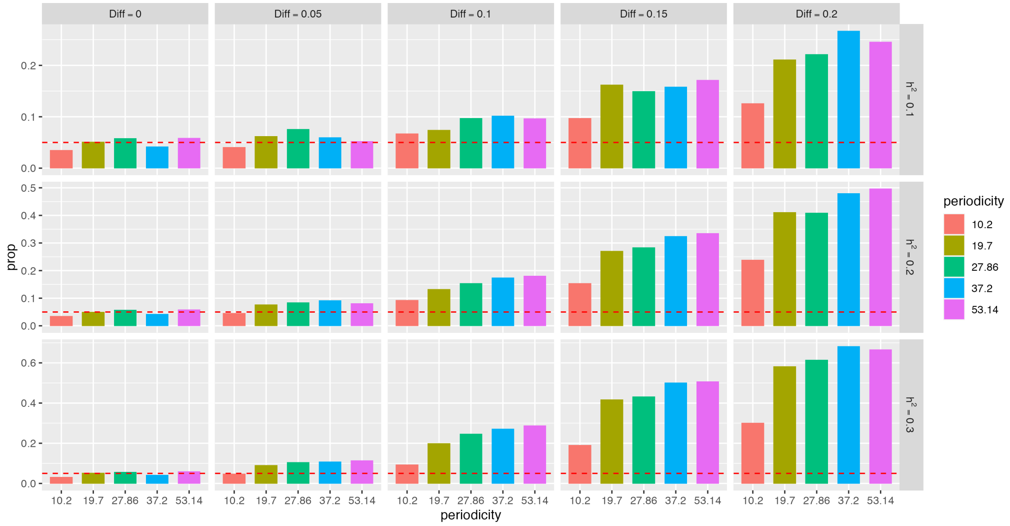


Figure 31: Bar plots for the power of spatialDEG with the estimated kernel. The ground truth is the Cosine kernel. The x axis provides the periodicity parameters and the y axis provides the proportion (prop) of significant genes. Diff denotes the difference in the means of the spatial components between the two datasets. The spot location is based on HFD1. The number of spots is $n_1 = n_2 = 1328$.

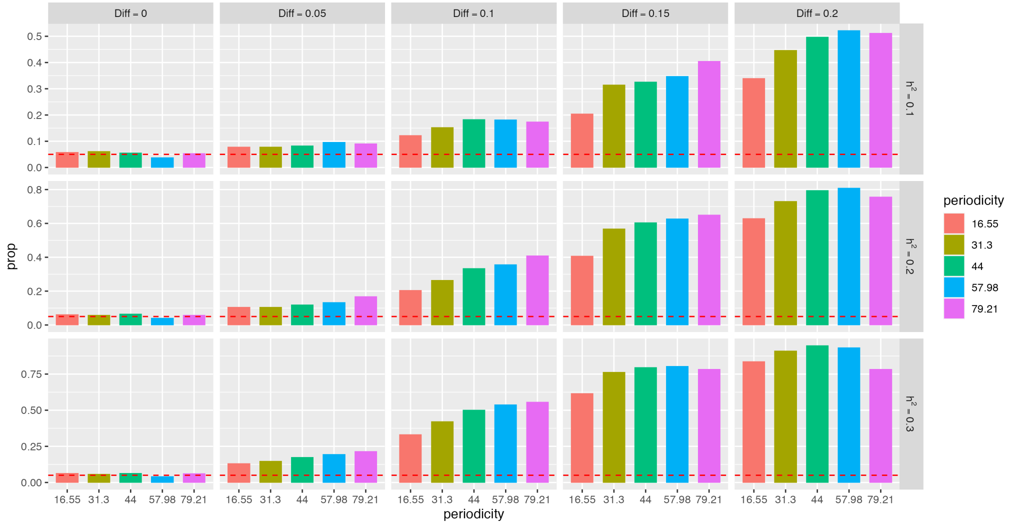


Figure 32: Bar plots for the power of spatialDEG with the estimated kernel. The ground truth is the Cosine kernel. The x axis provides the periodicity parameters and the y axis provides the proportion (prop) of significant genes. Diff denotes the difference in the means of the spatial components between the two datasets. The spot location is based on WT2. The number of spots is $n_1 = n_2 = 3613$.

### 3 Real data analysis

#### 3.1 Preprocessing of datasets

We first applied the pipeline Space Ranger from the 10x Genomics official website to the the Visium ST [12] data using the GRCh38 human reference genome or the mm10 mouse reference genome, also from the 10x Genomics official website, to generate spatial gene expression matrices. For both datasets, human dorsolateral prefrontal cortex (DLPFC) and mouse whole liver (WL), we log-transformed the raw count matrix using library size [5, 7]. We chose the 2000 most highly variable genes for each sample and performed the principal component (PC) analysis to obtain the top 15 PCs as the covariates while incorporating cell type information. We chose the top 20000 highly variable genes for each sample for the following spatialDEG analyses.

#### 3.2 ST datasets

##### 3.2.1 Human dorsolateral prefrontal cortex

For DLPFC, Maynard [6] used the pipelines Spatial Ranger for 10x Genomics Visium platform to generate feature-spot matrices in the form of h5 files and high-resolution and low-resolution images of tissues that were stored on the website for the program spatialLIBD. In their study, they profiled the total spatial gene expression in human postmortem DLPFC tissue sections of 12 samples from three donors. In summary, they obtained a median depth of 291 M reads for each sample, a mean of 3462 unique molecular indices, and a mean of 1734 genes per spot.

##### 3.2.2 Mouse whole liver

For WL, we also used the same pipelines Spatial Ranger for 10x Genomics Visium platform to generate feature-spot matrices in the form of h5 files and high-resolution and low-resolution images of the mouse WL tissue. For samples of mouse WL were included, two samples of which were from mice provided a high-fat diet (HFD), named HFD1 and HFD3, and two samples, named as WT1 and WT2, were from wild type (WT) mice. Similarly, we also profiled the spatial transcriptomics of mouse WL tissue sections from four samples using the 10x Genomics Visium platform. In summary, they have a mean depth of 263 M reads for four samples, as well as a mean 2053 unique molecular indices and a mean 2920 genes per spot.

##### 3.3 Differential analysis of DLPFC using spatialDEG

To evaluate the performance of spatialDEG when the samples were from the same specific condition, we conducted spatialDEG analysis on the four DLPFC samples 151507, 151508, 151509, and 151510, which were from the same neurotypical adult donors [6]. We first normalized these four samples as described above and performed the common dimension reduction method PCA on the data, thus we were able to obtain the top 15 PCs that incorporated cell type information for each sample. To avoid the effect of the cell type information within each sample, we regressed their PCs using multivariate linear regression for each sample and treated the residuals from the regression as the new gene expression matrix. Using pairwise combinations of these four samples with the cell type information removed, we could construct six pairs, referred to as 151507 vs 151508, 151507 vs 151509, 151507 vs 151510, 151508 vs 151509, 151508 vs 15150, and 151509 vs 151510, for analysis with spatialDEG. Specifically, we removed genes with less information that decreased the credibility of the analytical results. We also removed genes with summation of counts of less than 10 for each pair. After these pre-processing steps, we applied spatialDEG with the number of length scale and periodicity parameters $k_1 = k_2 = 10$ to analyze these six pairs. With the results for the six pairs generated by spatialDEG, we first had to remove genes that were inconsistent in their selection of the types of kernels between the null hypothesis and alternative hypothesis, such as Gaussian kernel for null hypothesis but Cosine kernel for alternative hypothesis, or vise versa, because the differences in kernels would lead to a greater difference in the log-likelihoood. The computational times for the six pairs 151507 vs 151508, 151507 vs 151509, 151507 vs 151510, 151508 vs 151509, 151508 vs 15150, and 151509 vs 151510 were approximately 7.5 h, 5.4 h, 5.8 h, 6.2 h, 5.9 h, and 12.7 h, respectively. All the real data were analyzed on the Linux platform with a 2.1 GHz Intel Xeon Gold 6230 CPU and 3.8 GB RAM on 20 cores.

##### 3.4 Differential analysis of WL using spatialDEG

To evaluate the performance of spatialDEG when the samples were from different conditions, we applied spatialDEG to the real dataset mouse WL. Similar to the DLPFC samples, we also needed to normalize all samples, calculate the corresponding PCs, and regress the PCs using multivariate linear regression. Then, we constructed four pairs, referred to as HFD1 vs WT1, HFD1 vs WT2, HFD3 vs WT1, and HFD3 vs WT2. Note that the two samples in each pair were from different conditions. Next, we removed genes with a summation of count of less than 10 from each pair to avoid inaccurate estimations due to low counts. Finally, we performed analysis using our proposed method spatialDEG with the number of length scale and periodicity $k_1 = k_2 = 10$ on these four sample pairs gene by gene. With the analysis results from spatialDEG, we first removed genes that showed inconsistency in the selection of the type of kernel between the null hypothesis and alternative hypothesis. The computational times for these four pairs are listed in Supplementary Table 4.

#### 3.5 qq-plots of p-values for the six pairs of DLPFC samples

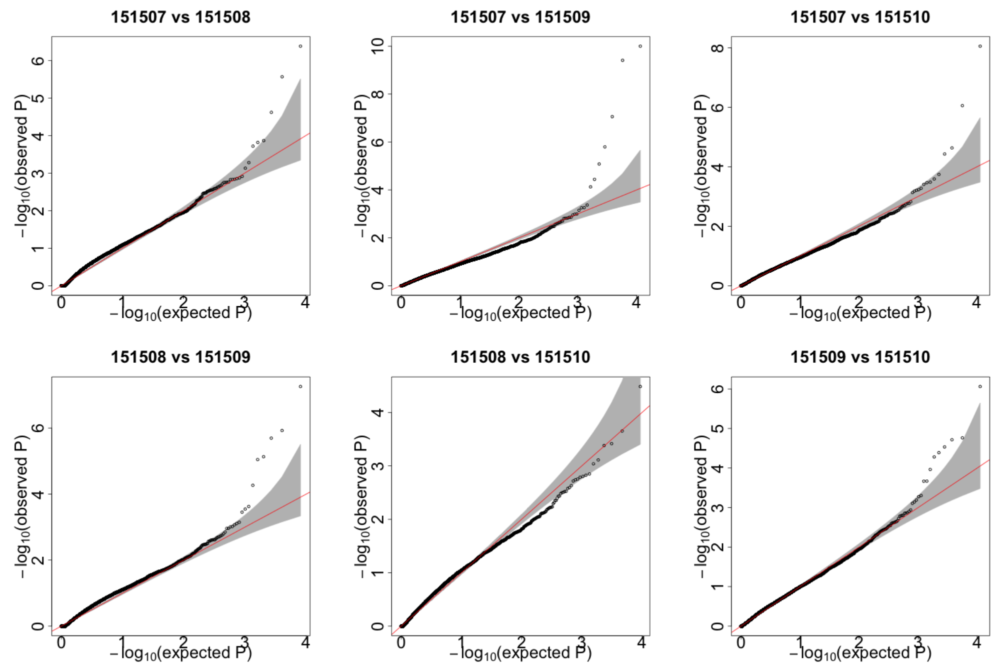


Figure 33: qq-plots of p-values generated by spatialDEG for the six pairs 151507 vs 151508, 151507 vs 151509, 151507 vs 151510, 151508 vs 151509, 151508 vs 15150, and 151509 vs 151510 from the real dataset DLPFC.

##### 3.6 DEGs of the four pairs of whole-liver samples

| pairs | Significant | Publication2021 | Publication2018 | time (h) |
|---|---|---|---|---|
| HFD1-WT1 | 33 | 10 | 8 | 2.6 |
| HFD1-WT2 | 66 | 17 | 28 | 4.6 |
| HFD3-WT1 | 40 | 16 | 15 | 2.7 |
| HFD3-WT2 | 86 | 27 | 34 | 6.4 |

Table 4: Number of significant genes identified by spatialDEG is shown in the Significant column. The number of significant genes reported in the previous publications [8] and [3] are in the columns Publication2021 and Publication2018. The computational times for the four pairs are shown in the column "time" (h)

##### 3.7 Pathway analysis of four pairs of whole-liver sample

We further conducted functional enrichment analysis of significant genes with a log fold change greater than 0.2 using gene ontology (GO); bubble plots for the four pairs are provided in Supplementary Figure 34. For all pairs, the FDR was set to 0.05. For HFD1 vs WT1, a total of 122 biological process (BP), 10 cellular component (CC), and 47 molecular function (MF) ontologies were enriched in significant genes identified by spatialDEG at an FDR of 0.05. Specifically, the four most significant BP ontologies, small molecule metabolic processes, lipid metabolic processes, carboxylic acid metabolic processes, and oxoacid metabolic processes are all different types of metabolic processes that are closely related to the consumption of a HFD [9]. Similarly, for HFD3 vs WT1, there were a total of 178 BP, 15 CC, and 53 MF ontologies enriched in significant genes. The top four BP ontologies were carboxylic acid metabolic processes, oxoacid metabolic processes, small molecule metabolic processes, and organic acid metabolic processes, which are also all metabolic processes that are altered under the condition of a HFD [2]. In addition, for HFD3 vs WT2, a total of 86 BP, 4 CC, 20 MF ontologies were enriched at an FDR of 0.05. In contrast to the above three pairs, no ontologies were enriched in significant genes identified by spatialDEG at an FDR of 0.05 for the pair HFD1 vs WT2.

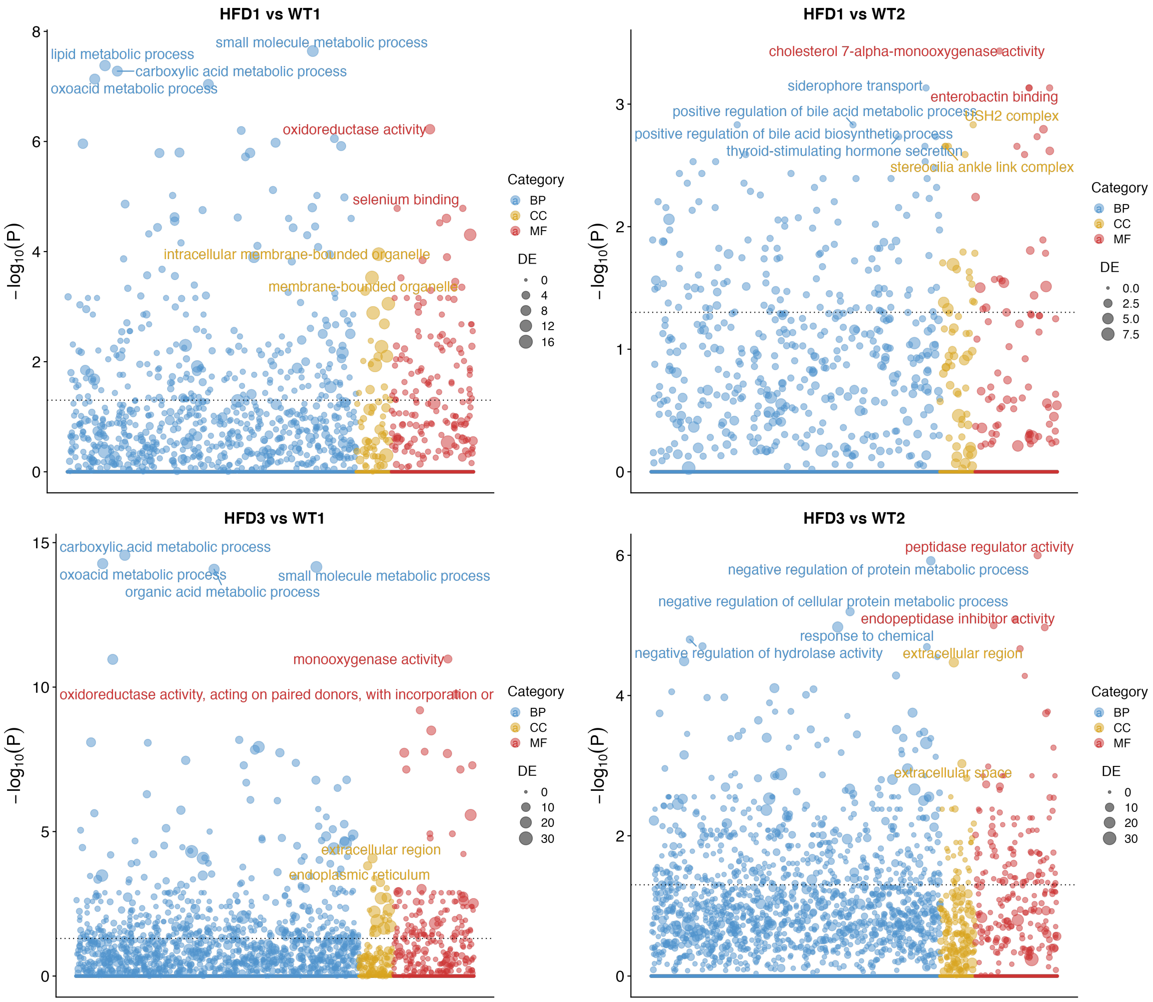


Figure 34: Bubble plots of pathway enrichment analysis for 17, 9, 34, and 35 differentially expressed genes for the pairs HFD1 vs WT1, HFD1 vs WT2, HFD3 vs WT1, and HFD3 vs WT2, respectively. The dashed line represents a p-value cutoff of 0.05. Gene sets are colored by category: GO biological processes (BP, blue), and GO cellular components (CC, yellow), GO molecular functions (MF, red).
